# Innate antiviral readiness drives the expansion of protective T stem cell memory against influenza

**DOI:** 10.64898/2025.12.06.692757

**Authors:** Ivan Tomic, David Ahern, Jeffrey Tomalka, Alba Escalera, Teresa Aydillo, Matthew D. Pauly, Nadine Rouphael, Seema S. Lakdawala, Naseem Sadek, Elizabeth Clutterbuck, Nisha Singh, Parvinder Aley, Hannah Robinson, Spyridoula Marinou, Stephanie P. Hao, Matthew Fish, Helder Nakaya, CHIM Study Group, Claudia Monaco, Adolfo Garcia-Sastre, Andrew J Pollard, Adriana Tomic

**Author notes:** Correspondence to: Adriana Tomic, Boston University. Equal contribution (Andrew J Pollard, Adriana Tomic).

## Abstract

The development of T-cell-based influenza vaccines relies on eliciting broad CD8+ T-cell immunity, wherein T stem cell-like memory (T_SCM_) cells serve as the ultimate long-lived reservoir for immune memory, thereby unlocking the potential for durable protection against viral drift and shift. However, the specific immunological cues that drive the robust expansion and functional preservation of this self-renewing, multipotent subset remain unknown. Here, utilizing multi-omic systems immunology in a pediatric cohort immunized with live attenuated influenza vaccine, we identified the determinants governing the expansion of influenza virus-reactive T_SCM_ cells. We show that a pre-existing state of innate antiviral readiness, defined by a plasmacytoid dendritic cell-associated type I interferon signature, is the requisite condition for a robust T_SCM_ expansion. Mechanistically, this baseline innate state enhances antigen priming and enforces a qualitative divergence in T-cell fate, driving responders toward a functionally poised, Th1-dominant phenotype while non-responders default to a dysfunctional, hyper-proliferative state. To determine the clinical relevance of this cellular subset, we analyzed an independent controlled human influenza challenge study. This validation revealed a critical functional division of labor in host defense: whereas pre-existing antibodies primarily mitigated symptom severity, the baseline frequency of influenza virus-reactive T_SCM_ cells was the strongest predictor of rapid viral load clearance. These findings establish that the expansion of durable cellular memory is not stochastic but is predetermined by the innate cytokine environment, providing a predictive biomarker for patient stratification and a validated target for adjuvants designed to expand the T_SCM_ reservoir deliberately.

## Introduction

Influenza remains a persistent global health threat due to the transient nature of vaccine-induced immunity and the continuous antigenic evolution of the virus (*1, 2*). Current vaccination strategies prioritize the induction of neutralizing antibodies against surface glycoproteins; while effective against matched strains, this protection is often short-lived and narrowly focused, leaving populations vulnerable to drifted seasonal variants and novel pandemic subtypes (*1*). Currently, several vaccine strategies to elicit durable, broadly reactive protective antibodies targeting conserved epitopes are being explored (*3, 4*). An alternative but not mutually exclusive approach to achieve durable, broad-spectrum protection consists of the induction of robust and long-lived cellular immunity targeting conserved internal viral proteins (*5*). In this context, T stem cell-like memory (T_SCM_) cells represent the long-sought goal of a T-cell-based vaccine design (*6–9*). Occupying the apex of the memory hierarchy, T_SCM_ cells possess the unique capacity for self-renewal and multipotency, serving as a sustainable reservoir that replenishes the effector pool upon antigen re-encounter and providing the cellular foundation for lifelong immunity (*10–13*).

Despite the recognized potential of T_SCM_ cells, translating this concept into clinical reality has stalled due to a fundamental knowledge gap: we lack the precise immunological requirements to reliably expand and maintain them. While modern vaccinology has excelled at maximizing the magnitude of short-lived effector responses (*14–18*), the specific signals that support the proliferation of stem-like memory without driving terminal differentiation are poorly understood. Furthermore, human vaccine responses exhibit significant inter-individual variability (*19*); it is currently unknown why some individuals mount a robust, functionally competent T_SCM_ recall response while others generate only abortive or exhausted memory populations. Unlocking the rules that govern this delicate balance, specifically, identifying the baseline determinants that support T_SCM_ amplification, is the critical prerequisite for engineering the next generation of universal vaccines.

To address these determinants, we employed a systems immunology approach in a unique cohort of children receiving the live attenuated influenza vaccine (LAIV). We selected this population because the adult immune system is heavily scarred by decades of cumulative exposure to influenza (*20, 21*), creating high background noise that obscures the signals of productive memory maintenance. Children, with their less skewed T-cell repertoires, provide a cleaner window into the foundational principles of immune boosting and memory persistence. Furthermore, as a replication-competent mucosal vaccine, LAIV mimics the natural kinetics of viral infection, allowing us to dissect the interplay between innate sensing and adaptive memory formation in a clinically relevant context (*22–24*).

In this study, we define the immunological rules of engagement for T_SCM_ expansion. We demonstrate that the capacity to expand a robust T_SCM_ pool is predetermined by a specific baseline state of innate antiviral readiness, driven by a plasmacytoid dendritic cell (pDC)-associated type I interferon (IFN) signature. We show that this innate alertness is not merely a biomarker but a functional driver that enhances antigen presentation and dictates the qualitative quality of the memory response, favoring a poised, Th1-dominant phenotype over a dysfunctional proliferative state. Finally, we bridge the gap between cellular phenotype and clinical protection using a controlled human influenza challenge (CHIM) model. Our analysis uncovers a distinct functional dichotomy in protective immunity, in which antibodies and T_SCM_ cells play separate but complementary roles in controlling symptoms versus clearing viral load. Collectively, these data provide a systems-level roadmap for targeting the innate-adaptive axis, offering a concrete strategy to improve vaccine durability by deliberately expanding the T_SCM_ reservoir.

## Results

### Prior LAIV exposure dictates T_SCM_ response magnitude

We investigated whether prior vaccination with a live attenuated influenza vaccine (LAIV) enhances the expansion of T_SCM,_ a critical population for durable immunity, in a pediatric cohort (n = 19, ages 4-6) (**Fig. 1A**). Given their rarity, the precise identification of T_SCM_ cells was crucial. We employed a stringent, multi-parameter cytometry strategy to define T_SCM_ cells (CD45RA⁺CCR7⁺CD95⁺CD27⁺CD28⁺) (**Fig. 1B**), distinguishing them from naïve (T_N_; CD95-), central memory (T_CM_; CD45RA-), and effector memory (T_EM_/T_EMRA_; CCR7-) subsets. Using this gating strategy, we found that T_SCM_ cells constituted approximately 1.5% of the total CD8+ T cell compartment at baseline (**Fig. 1B**), a frequency consistent with published estimates (*9*). Following a single LAIV dose, we observed significant inter-individual heterogeneity: 9 individuals showed an increase, while 10 individuals showed a decrease, or no change in T_SCM_ frequency (**Fig. 1E**).

**Figure 1.**
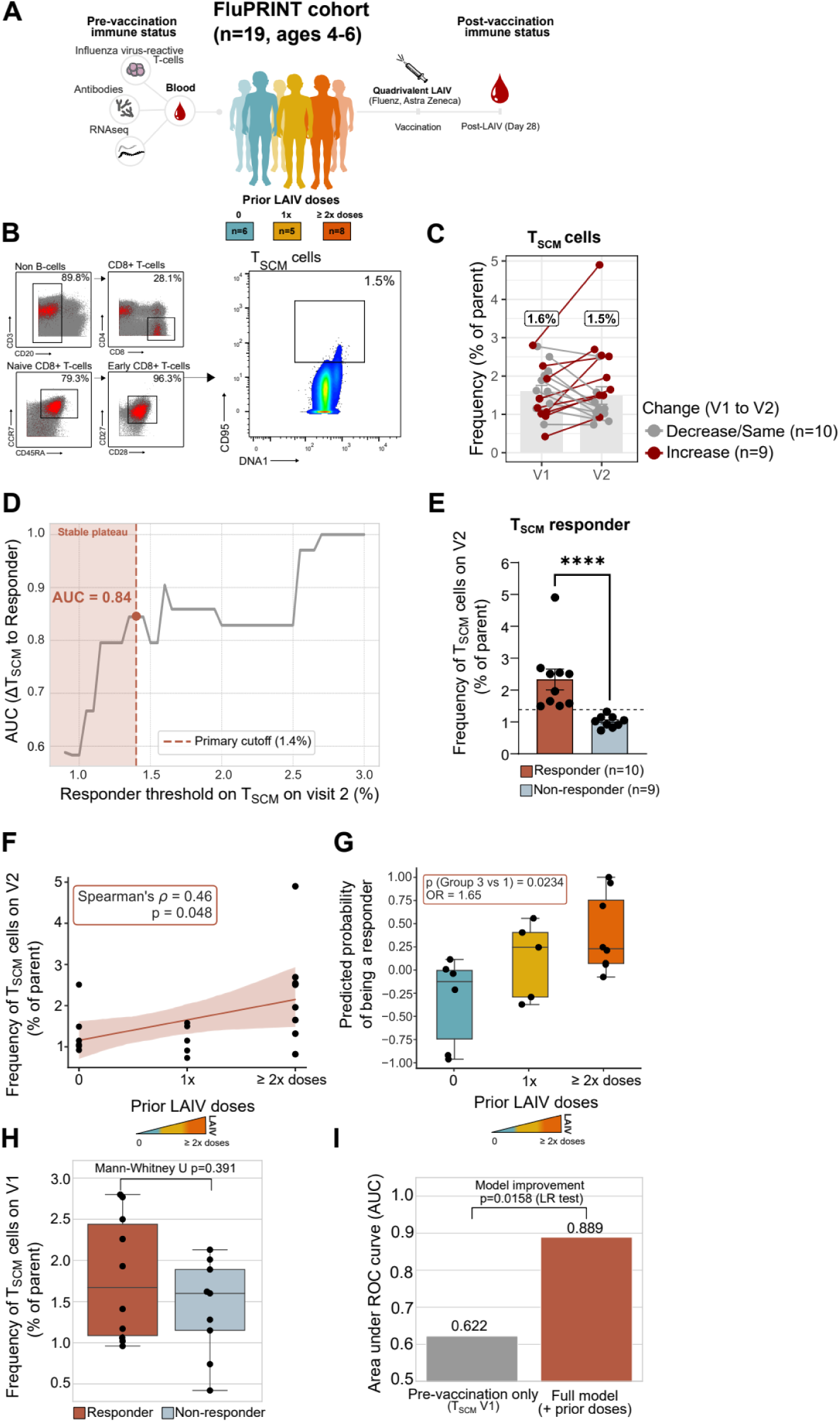
Study design and characterization of CD8+ T stem cell-like memory (T_SCM_) responses to LAIV in the FluPRINT cohort. **(A)** FluPRINT study overview showing the cohort of 19 children (ages 4-6 years) stratified by prior Live Attenuated Influenza Vaccine (LAIV) doses: 0 doses (n=6), 1 dose (n=5), or ≥2 doses (n=8), with immune assessment pre-vaccination and 28 days post-vaccination. **(B)** Mass cytometry gating strategy identifying CD8+ TSCM cells via sequential gating of CD3+, CD8+, naive (CD45RA+CCR7+), early (CD27+CD28+), and CD95+ markers. **(C)** Paired analysis of TSCM frequencies (% of parent) at Visit 1 (V1) and Visit 2 (V2) for subjects with increasing (red) or stable/decreasing (grey) responses. **(D)** Receiver operating characteristic analysis determining the responder threshold (1.4%) based on the stable plateau of the Area Under the Curve (AUC). **(E)** V2 TSCM frequencies stratified by responder and non-responder status relative to the 1.4% cutoff. **(F)** Correlation between V2 TSCM frequency and prior LAIV doses. **(G)** Predicted probability of being a responder stratified by prior LAIV exposure groups. **(H)** Comparison of baseline (V1) TSCM frequencies between responders and non-responders. **(I)** Comparison of predictive model performance (AUC) between pre-vaccination data only (grey) and the full model including prior doses (orange). Significance, denoted by specific p-values (e.g., p=0.048, p=0.0234), was computed using Spearman’s rank correlation for associations, Mann-Whitney U test for group differences, and the Likelihood Ratio test for model improvement comparisons. The odds ratio (OR) is indicated for probability assessments. Box plots represent the median with interquartile range and individual data points, while bar plots represent the mean with standard error. Shaded regions in correlation plots depict the 95% confidence interval.

To objectively classify the heterogeneous T_SCM_ response, we determined a responder threshold using a data-driven approach (**Materials and Methods**). We systematically tested a range of post-vaccination T_SCM_ frequencies (visit 2, V2) to identify the optimal cutoff that maximized the ability of the change in T_SCM_ cells (ΔT_SCM_) to discriminate responders. This analysis identified 1.4% as the optimal and most robust threshold (**Fig. 1D**). At this cutoff, ΔT_SCM_ served as an excellent classifier, achieving a high Area Under the Curve (AUC) of 0.844 in the receiver operating characteristic (ROC) curve (**Fig. 1D**). The validity of this 1.4% cutoff is underscored by its stability, as the high predictive power of ΔT_SCM_ remains consistent across a stable plateau of potential thresholds (**Fig. 1D**). At this threshold, we identified 10 responders (individuals with frequency of T_SCM_ following vaccination over 1.4%) and nine non-responders (**Fig. 1E**).

Using this metric, we found a clear dose-dependent relationship between prior LAIV immunization and the magnitude of the T_SCM_ response (**Fig. 1F**). We observed a statistically significant positive correlation between the number of prior LAIV doses and the post-vaccination T_SCM_ frequency (Spearman’s ρ = 0.46, p = 0.048) (**Fig. 1F**). Building on this, our multivariate logistic regression model confirmed that prior vaccination history was the key driver of the response. After adjusting for potential confounders, the number of prior LAIV doses remained a statistically significant predictor of responder status. As illustrated in **Figure 1G**, subjects with a history of two or more prior LAIV doses had a significantly higher predicted probability of being classified as a T_SCM_ responder compared to the vaccine-naïve group (p = 0.0234, Odds Ratio = 1.65). This finding remained robust even with the inclusion of additional covariates; the dose-response relationship remained significant after adjusting for baseline humoral immunity (HAI) (p = 0.0308, **Fig. S1**).

To critically validate that this dose-response was a direct consequence of vaccination history and not pre-existing immunity (i.e., subjects with more prior vaccinations having higher baseline T_SCM_ levels), we employed a systematic multivariate modeling approach. First, the pre-vaccination (V1) frequency of T_SCM_ cells showed no significant difference between responders and non-responders (Mann-Whitney U test, p = 0.391) (**Fig. 1H**). Next, we formally demonstrated that a logistic regression model including vaccination history (“*Full model*”) was a significantly better predictor of responder status (AUC = 0.889) than a model using only pre-vaccination TSCM levels (AUC = 0.622) (Likelihood-Ratio Test, p = 0.0158) (**Fig. 1I**).

Together, these data provide rigorous statistical evidence that prior LAIV vaccination primes the T_SCM_ cell compartment, leading to a more robust memory response upon subsequent immunization.

### Responders expand a functionally poised influenza virus-reactive T_SCM_ population

Having established that prior LAIV exposure dictates the quantitative magnitude of the T_SCM_ response (**Fig. 1**), our central objective was to define the qualitative nature of these cells and confirm their antigen specificity. An increase in cell numbers (our responder metric) reveals a capacity to respond. Still, it was crucial to determine whether this response was a successful, generating stable, functional influenza virus-reactive memory, or abortive, resulting in a dysfunctional, exhausted cell population.

To dissect this, we sought to define the precise functional state of the influenza virus-reactive cells within our LAIV cohort. Using mass cytometry, we employed a virtual sort to isolate *bona fide* influenza virus-reactive CD8+ T cells. Specifically, we identified cells upregulating activation-induced markers (AIM+), CD137, and CD69 upon influenza peptide stimulation and subtracted the background signal from matched unstimulated controls for each sample (**Materials & Methods**). This subtraction ensured that only genuinely influenza virus-reactive T cells were analyzed, yielding a total of 242,505 influenza virus-reactive CD8+ T cells from 38 samples for downstream analysis (**Fig. 2A**).

**Figure 2.**
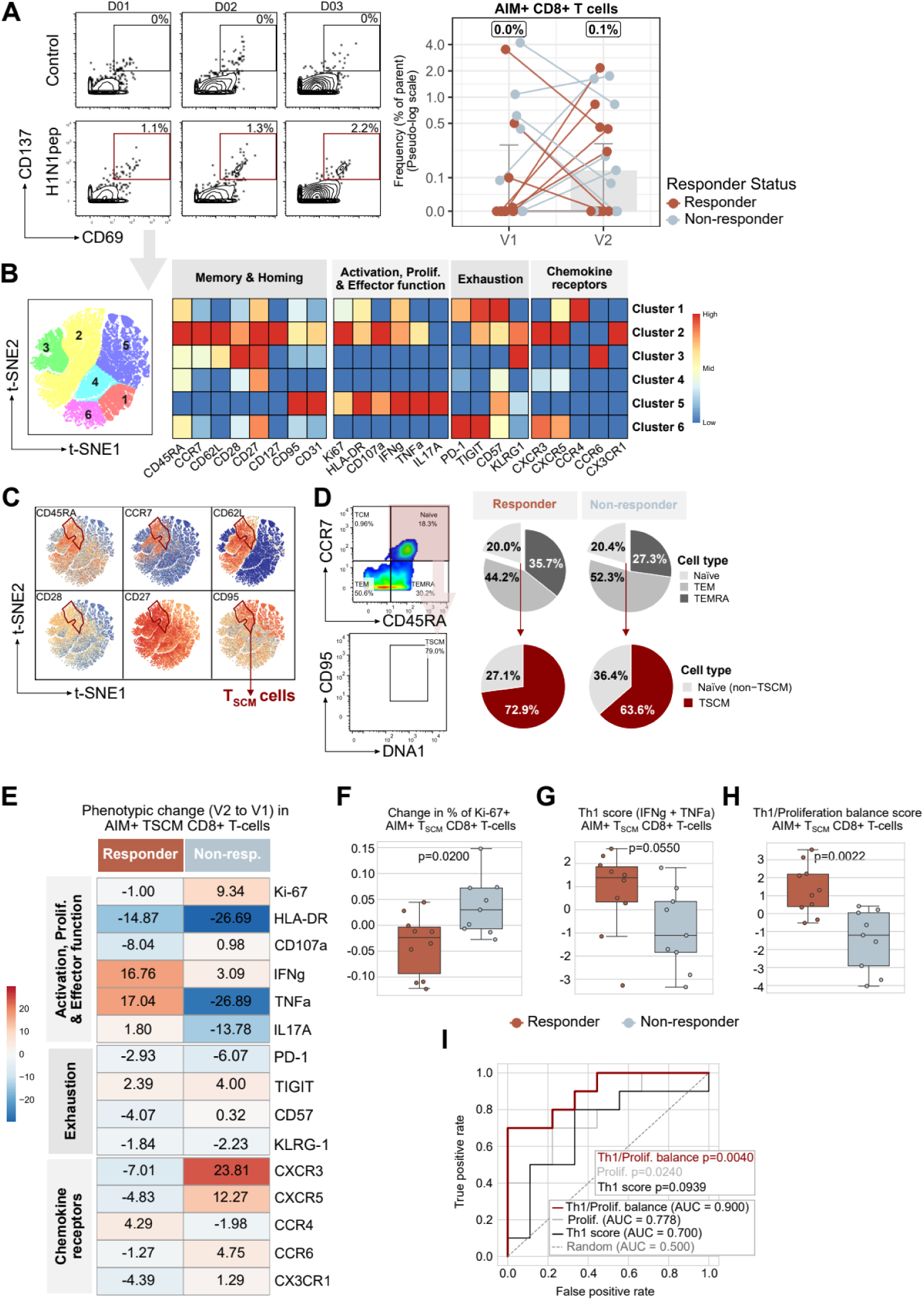
Phenotypic and functional characterization of influenza virus-reactive CD8+ T stem cell-like memory cells. **(A)** Identification of influenza virus-reactive T cells using an Activation Induced Marker (AIM) assay. Representative flow cytometry plots display CD137 and CD69 expression on CD8+ T cells following stimulation with H1N1 peptides (H1N1pep) or media control. The line graph (right) shows the frequency of AIM+ (CD69+CD137+) CD8+ T cells at Visit 1 (V1) and Visit 2 (V2) for responders (red) and non-responders (blue). **(B)** Unsupervised clustering of AIM+ CD8+ T cells. The heatmap (top) depicts the expression intensity of markers categorized by function (Memory & Homing, Activation/Proliferation/Effector function, Exhaustion, and Chemokine receptors) across 6 identified clusters, visualized spatially in the t-SNE plot (left). **(C)** t-SNE visualization of key marker expression (CD45RA, CCR7, CD62L, CD28, CD27, CD95) used to phenotypically identify TSCM cells. **(D)** Distribution of memory subsets within the AIM+ CD8+ T cell compartment. Top charts show the proportions of Naive, TEM, and TEMRA cells for responders and non-responders. Bottom charts detail the composition of the Naive-like compartment, distinguishing TSCM (red) from Naive (non-TSCM) cells (grey/black). **(E)** Heatmap showing the phenotypic change (V2 to V1) in AIM+ TSCM CD8+ T cells for responders and non-responders. Values and colors indicate the magnitude and direction (red for increase, blue for decrease) of change in expression for activation, proliferation, effector, exhaustion, and chemokine markers. **(F)** Boxplot comparing the change in proliferation (Ki-67+ frequency) within AIM+ TSCM cells between groups (p=0.0200). **(G)** Boxplot comparing the Th1 score (sum of IFNg and TNFa expression) within AIM+ TSCM cells (p=0.0550). **(H)** Boxplot comparing the Th1/Proliferation balance score (p=0.0022). **(I)** Receiver Operating Characteristic (ROC) analysis evaluating the performance of functional metrics in predicting responder status. Curves are shown for the Th1/Proliferation balance score (red, AUC=0.900, p=0.0040), Proliferation only (grey, AUC=0.778, p=0.0240), and Th1 score only (dotted, AUC=0.700, p=0.0939). Statistical significance was determined using Mann-Whitney U tests for group comparisons and likelihood ratio tests for model performance. Boxplots represent the median with interquartile range.

To map the phenotypic landscape of the response, we performed unsupervised clustering of all AIM+ CD8+ T-cells, identifying six distinct clusters (**Fig. 2B**, *left*). These clusters were defined by differential expression of markers for memory and homing, activation and function, exhaustion, and chemokine reception (**Fig. 2B**, *right*). This revealed that the clusters segregated along two primary axes: 1) a memory-phenotype axis, distinguishing stem cell memory-like (C2, C3) from effector memory (C4, C5, C6) and T_EMRA_-like (C1) populations, and 2) a functional axis, separating quiescent clusters (C3, C4, C6) from highly active, polyfunctional, and proliferative clusters (C1, C2, C5). While this global analysis revealed the complexity of the landscape, it did not reveal significant differences in frequency between responders and non-responders (**Fig. S2**). This suggested the critical differentiator was not which cluster cells belonged to, but rather a more subtle functional state change within a specific population. Given our hypothesis, we therefore focused our analysis on the AIM+ cells within the T_SCM_ compartment.

A focused t-SNE analysis identified the T_SCM_ population within cluster 2, characterized by expression of CD95, CD27, CD28, CCR7, CD45RA, and CD62L, as previously reported (*9*) (**Fig. 2C**, *red circle*). Manual cytometry gating further confirmed that most AIM+ T-cells have a T_EM_ phenotype (around 50%), 30% T_EMRA_, and 20% naïve phenotype, within which up to 70% have a T_SCM_ phenotype, without differences between responders and non-responders (**Fig. 2D**).

We next sought to define the qualitative differences in the AIM+ T_SCM_ compartment between responders and non-responders by analyzing the phenotypic change from V1 to V2 (**Fig. 2E**). This revealed a stark functional divergence. Responders were characterized by a functionally poised signature: a decrease in the proliferation marker Ki-67 (mean change: -1.00) and the activation marker HLA-DR (-14.87), coupled with a substantial increase in Th1 effector function markers IFNγ (+16.76) and TNFα (+17.04). Conversely, non-responders displayed a dysfunctional, proliferative profile, characterized by a robust rise in Ki-67 (+9.34) and HLA-DR (+26.69), but a loss of Th1 function, particularly in TNFα (-26.89).

This proliferative state was a key differentiator. A focused analysis confirmed that the change in Ki-67+ AIM+ T_SCM_ cells was significantly different between groups, with responders decreasing proliferation and non-responders increasing it (p=0.0200; **Fig. 2F**). This was directly linked to functional quality; responders displayed a higher Th1 score (IFNγ + TNFα (p=0.0550; **Fig. 2G**) and a significantly higher Th1/Proliferation balance score (p=0.0022; **Fig. 2H**).

Finally, a comparative ROC analysis confirmed that this Th1/Proliferation balance score was the most robust and accurate metric for classifying responders, achieving an Area Under the Curve (AUC) of 0.900. This outperformed the predictive power of the Th1 score alone (AUC = 0.700) or the proliferation marker alone (AUC = 0.778) (**Fig. 2I**). This demonstrates that a successful AIM+ T_SCM_ response is defined not just by expansion, but by a qualitative shift toward a functionally poised (Th1-high) and quiescent (Ki-67-low) state.

### A pre-existing signature of broad innate priming predicts the quantitative and qualitative fate of the T_SCM_ response to live attenuated influenza vaccination

Our data establishes that a successful T_SCM_ responder to LAIV is defined by both a significant quantitative expansion of the T_SCM_ compartment (**Fig. 1**) and a superior qualitative functional fate of those cells (**Fig. 2**). This raised the central question of our study: what molecular determinants, present at baseline, prime an individual to mount this functionally poised and numerically robust T_SCM_ response to LAIV? As a first exploratory step to identify this predictive signature, we interrogated the pre-vaccination whole-blood transcriptomes and plasma proteomics (O-Link) of T_SCM_ responders and non-responders (defined at day 28 post-LAIV).

Differential expression (DE) analysis revealed a distinct, pre-existing transcriptional dichotomy between the two groups. T_SCM_ responders displayed a marked and coherent upregulation of genes central to innate antiviral defense and the interferon (IFN) pathway (**Fig. 3A, B**). This signature was dominated by canonical interferon-stimulated genes (ISGs), including viral sensors (*19*) like *DDX58* (RIG-I), core components of IFN signal transduction (*25*) (*STAT1*, *STAT2*), and a broad array of canonical interferon-stimulated genes (ISGs), including influenza restriction factors *MX1* (*26*) and *RSAD2* (Viperin) (*27*), and *IFI44* (*28*) and *IFI44L* (*29*), all recognized for their antiviral action against the influenza virus (*30*) (pAdj < 0.05, Log2FC > 1) (**Fig. 3A, B**). In contrast, non-responders exhibited significant upregulation of genes associated with immune homeostasis and immune regulation, such as the transcription factors *BACH2* (*31*) and *FOXO1* (*32*) (pAdj < 0.05, Log2FC > 1) (**Fig. 3A, B**). Unsupervised hierarchical clustering of the top differentially expressed genes (DEGs) confirmed that they constitute a co-regulated transcriptional program, which robustly segregated responders from non-responders based on this baseline signature alone (**Fig. 3B**).

**Figure 3.**
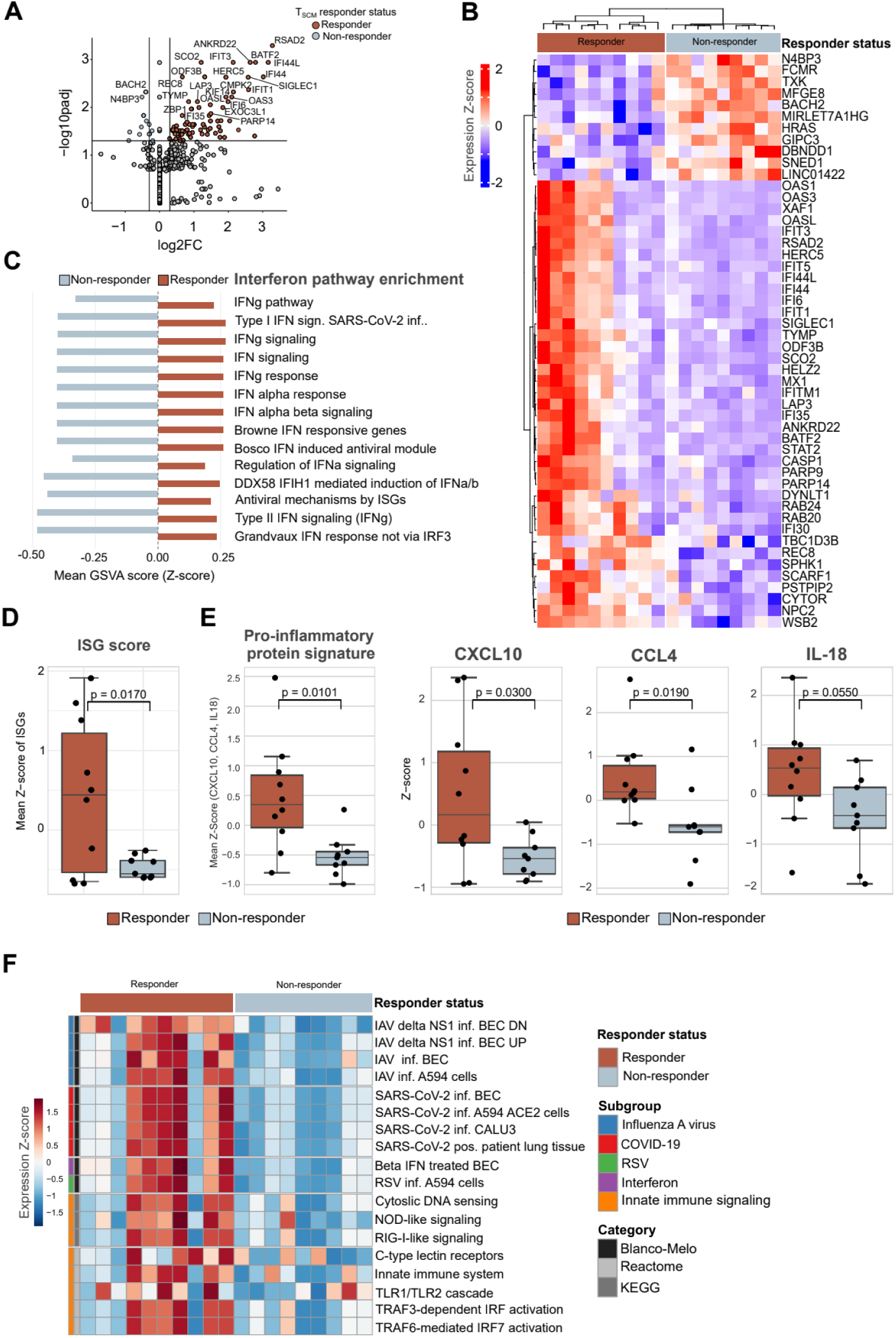
Transcriptional and proteomic signatures associated with TSCM responsiveness. **(A)** Volcano plot illustrating differential gene expression between responders (red) and non-responders (blue). The x-axis represents the log2 fold change (log2FC), and the y-axis represents the significance (-log10 adjusted p-value). Upregulated genes in responders, such as *RSAD2*, *IFI44L*, and *ANKRD22*, are highlighted in red. **(B)** Heatmap displaying the expression Z-scores of the top differentially expressed genes across individual subjects, stratified by responder (red bar) and non-responder (blue bar) status. **(C)** Bar plot comparing Interferon pathway enrichment between groups. Bars represent the Mean Gene Set Variation Analysis (GSVA) score (Z-score) for various interferon-related pathways in responders (red) and non-responders (blue). **(D)** Boxplot comparing the Interferon Stimulated Gene (ISG) score (calculated as the mean Z-score of ISGs) between responders (red) and non-responders (blue) (p=0.0170). **(E)** Analysis of the pro-inflammatory protein signature. Boxplots compare the composite Mean Z-score of three proteins (CXCL10, CCL4, IL-18) between groups (p=0.0101), followed by comparisons for individual proteins: CXCL10 (p=0.0300), CCL4 (p=0.0190), and IL-18 (p=0.0550). **(F)** Heatmap of gene set enrichment scores across subjects. Rows represent specific gene sets related to viral infections (Influenza A, COVID-19, RSV) and innate immune signaling (Interferon, RIG-I-like signaling, etc.), annotated by category (Blanco-Melo, Reactome, KEGG). Columns represent individual subjects colored by responder status. Red indicates upregulation and blue indicates downregulation of the respective gene sets. Statistical significance for boxplots was determined using Mann-Whitney U tests, with p-values indicated above the brackets. Boxplots represent the median with interquartile range.

To elucidate the functional coherence of this signature, we performed Gene Set Variation Analysis (GSVA) on a broad panel of interferon-related pathways. This analysis confirmed the DEG findings, revealing a systematic and significant enrichment (pAdj < 0.05; mean GSVA Z-score > 0) for nearly all IFN-related pathways in responders, including the IFNγ pathway and IFNα/β signaling, while non-responders showed a consistent negative enrichment (mean GSVA Z-score < 0) (**Fig. 3C**).

We next sought to quantify this innate readiness using integrated, multi-omic signatures. At the RNA level, a composite ISG score derived from the mean Z-score of 32 canonical ISGs was significantly elevated in responders (Welch’s Two Sample t-test, p = 0.0170; **Fig. 3D**). To determine if this transcriptional state translated to a functionally active, circulating proteome, we analyzed baseline plasma proteomics (O-link). This analysis revealed a striking validation: a pro-inflammatory protein signature, independently identified via Lasso regression as a robust predictor (CV-ROC AUC = 0.85; **Fig. S3A, B**) and comprised of CXCL10, CCL4, and IL18, was also statistically significant (Mann-Whitney U test, p=0.0101) and strongly upregulated in responders (Cohen’s d = 1.38; **Fig. 3E***, left panel*). Furthermore, the Lasso model also identified a distinct signature for non-responders, characterized by the specific upregulation of FGF-19 and IL-17C, a factors primarily involved in metabolic homeostasis and tissue repair (*33*) (**Fig. S3C**), further confirming that the responder/non-responder dichotomy is defined by active, opposing biological programs. This demonstrates a direct link between the baseline ISG transcript (**Fig. 3D**) and a pre-activated proteomic milieu (**Fig. 3E**).

Finally, to define the full mechanistic depth of this signature, we performed GSVA (pAdj < 0.05) on a broader panel of innate immune pathways from the Blanco-Melo, KEGG, and REACTOME databases (**Fig. 3F**). The Blanco-Melo dataset (*34*) provides a foundational benchmark, defining the precise ISG-driven transcriptional architectures of human epithelial cells during active infection with respiratory viruses (e.g., influenza A virus (IAV), severe acute respiratory syndrome coronavirus 2 (SARS-CoV-2), and respiratory syncytial virus (RSV)). This analysis revealed potent, uniform enrichment (*red*) across a broad spectrum of viral response signatures (IAV, SARS-CoV-2, RSV), suggesting that responders possess a broad, non-specific antiviral readiness state (**Fig. 3F**). Furthermore, this enrichment was most substantial for pathways representing a maximal, IFN-driven state, such as infection with IAV delta-NS1 (a hyper-IFN-inducing mutant (*35*)) and direct IFNβ treatment. Crucially, this state of antiviral readiness was not restricted to downstream ISGs. The analysis confirmed coordinate enrichment of upstream pathogen-sensing modules from KEGG and REACTOME, including cytosolic DNA sensing, RIG-I-like signaling, NOD-like signaling, C-type lectin receptors, and TLR1/TLR2 cascades (**Fig. 3F**). This demonstrates that the predictive signature in T_SCM_ responders is a deeply coherent state of heightened preparedness, spanning the entire innate immune axis from initial pathogen sensing to terminal effector gene expression.

Collectively, this exploratory multi-omic analysis validated a model: a successful T_SCM_ response is predicted by a baseline state of broad innate readiness, quantifiable from the initial sensor transcript (**Fig. 3D**) to the final, circulating effector protein (**Fig. 3E**).

### A pDC-associated cascade governs T_SCM_ response to LAIV via a dual-path mechanism

Our multi-omic analysis has established that a successful LAIV response, defined by a quantitative T_SCM_ expansion of a qualitatively functionally poised nature, is predetermined by a pre-existing IFN signature. This raised the critical, mechanistic question: how does this innate signature translate to a superior adaptive response? To transition from a correlative theme to a causal hypothesis, we therefore sought to develop a system-level causal model to define the pathway’s structure and cellular origin. In canonical immunity, type I and II IFNs are known to bridge this gap by driving the maturation, antigen processing, and cross-presentation capabilities of antigen-presenting cells (APCs) (*36–39*). We therefore hypothesized that our baseline ISG signature would be linked to enhanced antigen presentation. Testing this, we returned to our baseline transcriptomics and confirmed this hypothesis: T_SCM_ responders showed significant baseline enrichment (FDR < 0.05) for pathways including class I MHC-mediated antigen processing and presentation and antigen processing via cross-presentation (**Fig. 4A**), suggesting the pre-existing state in responders is not just passively antiviral but is actively poised to enhance T-cell priming.

**Figure 4.**
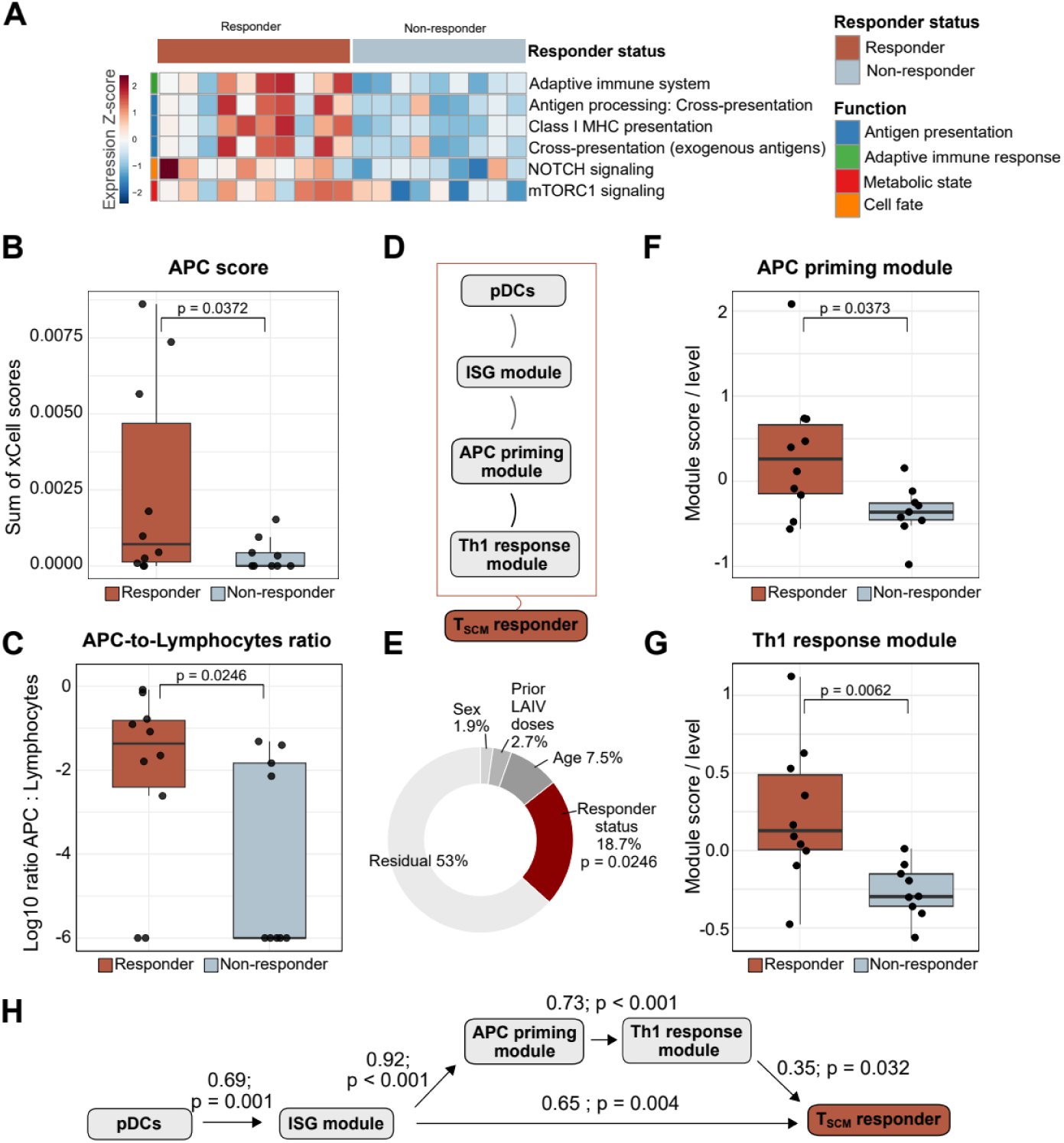
Modular analysis of the innate and adaptive immune drivers of TSCM responses. **(A)** Heatmap displaying the expression Z-scores of differentially enriched gene sets between responders (red) and non-responders (blue). Gene sets are annotated by functional categories: Antigen presentation (blue), Adaptive immune response (green), Metabolic state (pink), and Cell fate (orange). **(B)** Boxplot comparing the aggregate Antigen Presenting Cell (APC) score (sum of xCell enrichment scores for pDCs and macrophages) between responders (red) and non-responders (blue) (p=0.0372). **(C)** Boxplot comparing the log10 ratio of APCs to lymphocytes (T- and B-cells) between groups (p=0.0246). **(D)** Schematic diagram proposing the sequential relationship between modules: pDCs activate the ISG module, leading to the APC priming module, the Th1 response module, and ultimately the TSCM responder phenotype. **(E)** Variance partition analysis (donut chart) showing the percentage of variance in the TSCM response explained by responder status (18.7%, p=0.0246), relative to demographic factors (sex, age) and prior LAIV exposure. **(F)** Boxplot comparing the expression levels of the APC priming module between responders (red) and non-responders (blue) (p=0.0373). **(G)** Boxplot comparing the expression levels of the Th1 response module based on responder status (p=0.0062). **(H)** Path analysis/Structural Equation Modeling (SEM) quantifying the strength and significance of the relationships between the identified modules. Arrows indicate the direction of influence, with numbers representing path coefficients and p-values indicating statistical significance.

This enrichment of APC pathways suggested a heightened functional state, and we reasoned that the APCs themselves might drive it. To investigate the cellular basis for this dual IFN and APC signature, we performed computational deconvolution of the whole-blood transcriptomes. This analysis confirmed our hypothesis, revealing that a composite baseline signature of pDCs and macrophages was significantly elevated in responders (Wilcoxon p = 0.037, **Fig. 4B**). Furthermore, this translated to a significant shift in immune architecture, with responders showing a higher baseline ratio of APCs (pDCs + Macrophages) to total lymphocytes (T + B-cells) (Wilcoxon p = 0.0246, **Fig. 4C**). This finding was specific, as individual APC populations like pDCs (p = 0.085) or macrophages (p = 0.103) alone, other non-APC lymphoid and myeloid populations (**Fig. S4A**), or a broader composite of all APCs were not significantly different (pDCs, cDCs and macrophages, p = 0.079; **Fig. S4B**).

While our deconvolution analysis identified a composite enrichment of both pDCs and macrophages in responders, determining the specific cellular initiator of the observed IFN signature is critical. To resolve this and model the complete pathway, we constructed a Structural Equation Model (SEM), a multivariate technique for testing and evaluating complex causal relationships (*40*). The critical objective of this modeling was to empirically resolve the cellular source of the IFN signature, given that our initial analysis identified a composite enrichment of both pDCs and macrophages. Although macrophages are abundant and capable of producing type I IFN, pDCs are the canonical primary producers (*39, 41–43*). To test this, we evaluated two competing causal architectures: one driven by baseline pDCs and an alternative driven by baseline macrophages. We hypothesized a complete, systems-level cascade: a high baseline pDC or macrophage potential leads to a robust early ISG score, which in turn drives APC maturation and presentation, leading to increased priming, culminating in a superior Th1 response and, ultimately, more robust expansion of the T_SCM_ population observed in responders (**Fig. 4D**). First, we defined these four *a priori* biological scores as composite variables (GSVA scores) derived from functionally coherent sets of biological features (see **Materials & Methods**): (1) pDC score, a calculated deconvolution score; (2) ISG score, a composite score computed as the mean of 32 scaled canonical ISG genes; (3) APC priming score, a composite score computed from five scaled genes (e.g., *FCGR1A*, *LAMP3*) and eight scaled protein measurements (e.g., CCL3, CD40); and (4) Th1 response score, a composite score computed from two scaled genes (*STAT1*, *TNFSF10*), five scaled proteins (e.g., CXCL9, IFNγ) and six scaled cell cluster frequencies.

We first sought to validate our responder status as a primary, biologically significant endpoint for modeling. A PERMANOVA (permutational multivariate analysis of variance) on the global, multi-omic signature data from all subjects (**Fig. 4E**) confirmed that responder status emerged as a highly significant driver of global variance (R2 = 0.1870, p = 0.0246), explaining 18.7% of the total variation in the dataset. In contrast, potential demographic confounders, such as sex (R² = 0.0185, p = 0.6986) and age (R² = 0.0747, p = 0.4888), were not significant, confirming responder status as a robust biological endpoint.

We next investigated the relationships between the key components of our hypothesized causal chain (**Fig. 4F, G**). Univariate comparisons confirmed that the downstream scores for the APC priming score (p = 0.0373; FDR = 0.0497; **Fig. 4F**) and Th1 response score (p = 0.0062; FDR = 0.0249; **Fig. 4G**) were all significantly elevated in responders. Critically, however, the hypothesized initiator of the cascade, the baseline pDC (p = 0.085) or macrophage (p = 0.120) score, failed to reach significance in this direct comparison (**Fig. S4A**). This suggested a more complex, multi-step relationship in which the effect of the baseline state was nonlinear. Indeed, this was confirmed by simple pairwise correlations and formal causal mediation analyses (**Table S1**), which failed to yield significant mediation effects (ACME p = 0.21, 0.843, 0.39). This underscored the inadequacy of simple linear models and the necessity of a systems-level approach.

The pDC-initiated SEM model demonstrated an excellent fit to our data (CFI = 0.992, RMSEA = 0.041; **Table S2**) and revealed a robust causal chain. It established that baseline pDCs, despite being non-significant in univariate tests, were a highly significant positive driver of the ISG score (Estimate = 237.259, z = 3.433, p = 0.001). This ISG score then significantly predicted the APC priming score (Estimate = 0.686, z = 3.873, p < 0.001), which in turn was a powerful predictor of the final Th1 response score (Estimate = 0.486, z = 5.005, p < 0.001). Most critically, the pDC model revealed a novel dual-path mechanism predicting the outcome. Responder status was significantly predicted by the terminal Th1 response score (Estimate = 1.039, z = 2.144, p = 0.032) and by a direct, independent shortcut path from the early ISG score (Estimate = 0.949, z = 2.902, p = 0.004). This dual-path model was highly explanatory, accounting for 85.4% of the total variance in responder status (R-Square = 0.854). In contrast, when we tested the alternative macrophage-initiated model, the mechanistic architecture proved inferior. While baseline macrophages could predict the ISG score (p = 0.003), the model explained less variance in the clinical outcome (R2 = 0.730), and the critical rapid response shortcut path, the direct link from ISG score to responder status, failed to reach significance (p = 0.717) (**Table S3**).

In conclusion, our findings establish that a pDC-initiated baseline state, rather than a generalized macrophage signature, predetermines vaccine response by branching into two key pathways: a direct, rapid interferon activation and a complete, stepwise cascade that primes the adaptive T-cell response.

### Pre-existing hyper-interferon state defines baseline immunotypes that predict vaccine-induced T_SCM_-response

The SEM provides a robust statistical causal framework, but it connects the outcome to an abstract ISG score as the central, causal hub. While this established a causal linkage, it did not identify the specific biological drivers. This raises two critical questions: 1) What is the precise, biological identity of the ISG score hub? 2) Do the statistical arrows of our SEM (e.g., from ISG score to APC priming score) represent known, physical, molecular interactions? To answer these questions and provide a final synthesis of our findings, we designed a three-step, sequential validation analysis.

First, to define the specific biological identity of the ISG score latent variable, we moved from the broad exploratory analysis of GSVA (**Figure 3)** to a supervised, discriminant analysis. We employed Partial Least Squares Discriminant Analysis (PLS-DA), a method ideal for feature selection in complex, collinear multi-omic datasets. Unlike the broad theme-finding of GSVA, PLS-DA is a supervised method that leverages the known responder labels to identify the minimal, non-redundant set of features with the maximal power to discriminate between the two groups. The PLS-DA confirmed this hypothesis (**Fig. 5A**). The primary axis of separation (Component 1) was defined by a broad, coherent program of canonical interferon signatures, including IFNα response (FDR = 0.027), IFNγ response (FDR = 0.027), and IFN signaling (FDR = 0.027) (**Fig. 5A**, *right panel*; **Table S4**). Critically, the single top-ranked discriminatory feature (FDR = 0.022) was a signature derived from cells infected with IAV lacking the IFN-antagonist protein NS1. Because the NS1 protein’s function is to suppress the host’s innate immune response (*35*), this delta-NS1 signature represents a massive, uninhibited hyper-interferon state. This provides a definitive biological definition for the ISG score in our mechanistic model.

**Figure 5.**
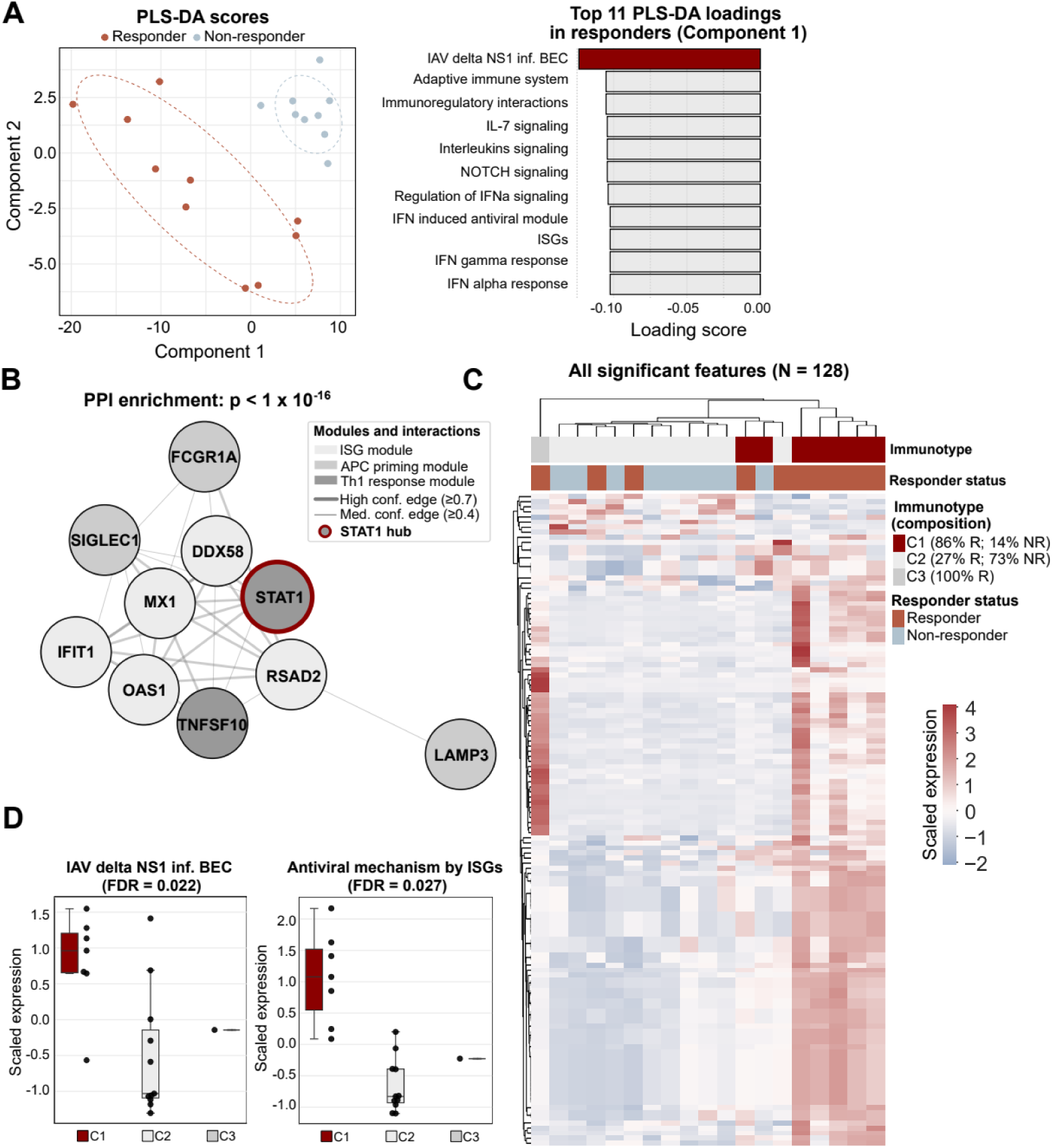
Multivariate modeling and identification of distinct immunotypes associated with LAIV response. **(A)** Partial Least Squares Discriminant Analysis (PLS-DA). The scores plot (left) visualizes the separation between responders (red) and non-responders (blue) along Component 1 and Component 2. The bar plot (right) lists the top 11 loading features contributing to Component 1, including influenza A virus (IAV) delta NS1 infection signatures and adaptive immune system pathways. **(B)** Protein-Protein Interaction (PPI) network analysis of significant features (p < 1 × 10⁻¹⁶). The network highlights a central STAT1 hub (red node) interacting with key antiviral genes such as *MX1*, *OAS1*, and *FCGR1A*. Edge thickness indicates confidence levels. **(C)** Unsupervised hierarchical clustering heatmap of all significant features (N=128). The analysis reveals three distinct clusters/immunotypes (C1, C2, C3) with varying proportions of responders (R) and non-responders (NR): C1 (86% R), C2 (27% R), and C3 (100% R). **(D)** Boxplots validating the functional differences between the identified immunotypes (C1, C2, C3) using scaled expression of IAV delta NS1 inf. BEC (FDR=0.022) and Antiviral mechanism by ISGs (FDR=0.027) gene sets. Boxplots represent the median with interquartile range.

Next, we sought to build the wiring diagram and physically validate the statistical arrows of our SEM (**Fig. 4H**). Our SEM hypothesized a statistical cascade from the ISG score to the APC priming score. We therefore performed a protein-protein interaction (PPI) network analysis to determine whether these top features constitute a random collection or a functionally integrated, physically cohesive system, as our model would predict. The analysis provided a definitive result: our features formed a densely interconnected network (p < 1.0 × 10^−16^) (**Fig. 5B**). Critically, the network was centered on the viral sensor DDX58 (RIG-I; retinoic acid-inducible gene I), a cytosolic pattern recognition receptor that mediates the induction of a type I IFN response (*44*), and the master IFN-inducible transcription factor, signal transducer and activator of transcription 1 (STAT1) (*45, 46*). Finding STAT1, the master ON switch for the IFN response (*46*), at the network’s hub is the ultimate validation. It is the physical transducer that receives the alarm signal (from DDX58) and, in turn, activates the genes that define our ISG score (e.g., MX1, OAS1). This network supported SEM findings, revealing direct, known interactions linking the ISG score molecules (MX1) to the APC priming molecules (LAMP3, FCGR1A) and, finally, to the Th1 response molecules (TNFSF10) (**Fig. 5B**).

Finally, to provide the ultimate translational validation for this entire framework, we asked: Is our responder classification, which we imposed based on a single parameter (**Fig. 1**), biologically real? We discarded our outcome labels and performed unsupervised consensus clustering on the entire baseline multi-omic dataset. This agnostic approach enables the data to reveal the underlying patient subgroups, or immunotypes, without any *a priori* assumptions. The clustering identified stable, pre-existing patient immunotypes (C1, n = 7; C2, n = 11) (**Fig. 5C**). These baseline immunotypes were strongly predictive of the ultimate vaccine response (p < 0.01). Subjects in immunotype C1 were predominantly responders (85.7%), whereas subjects in immunotype C2 were predominantly non-responders (72.7%) (**Fig. 5C**). Crucially, as shown in **Figure 5D**, these agnostically-defined immunotypes are defined by the identical biological signatures our PLS-DA identified. Subjects in the responder-like C1 immunotype showed significantly higher expression of our top two biomarker hits: the IAV delta NS1 infection signature (FDR = 0.022) and the antiviral mechanism by ISGs pathway (FDR = 0.027).

This finding provides a comprehensive, systems-level model: the immunotypes are the tangible, pre-existing individual states defined by the hyper-interferon signature, which in turn dictates the pDC potential that initiates the entire causal cascade, determining the success or failure of the T_SCM_ response to LAIV.

### T_SCM_ cells dictate viral control, not symptoms, in human influenza infection challenge

This raises the ultimate translational question: are the cells we identified, generated by childhood vaccination, capable of protecting against a subsequent infection? To determine if these cells, with their unique durability, can protect against an influenza virus exposure encountered later in adulthood, we analyzed a cohort of healthy adults (n = 9; ages 21-46) from a controlled human influenza challenge (CHIM) study (*47*) (**Fig. 6A**). Following baseline blood collection, subjects were infected intranasally with H3N2 (A/Perth/16/2009), and viral load and symptoms were monitored for 8 days while in quarantine (**Fig. 6A**). Baseline T-cell reactivity was assessed using AIM assay where PBMCs were stimulated with an entire influenza A virus peptide library exactly as for the FluPRINT cohort (**Materials & Methods**).

**Figure 6.**
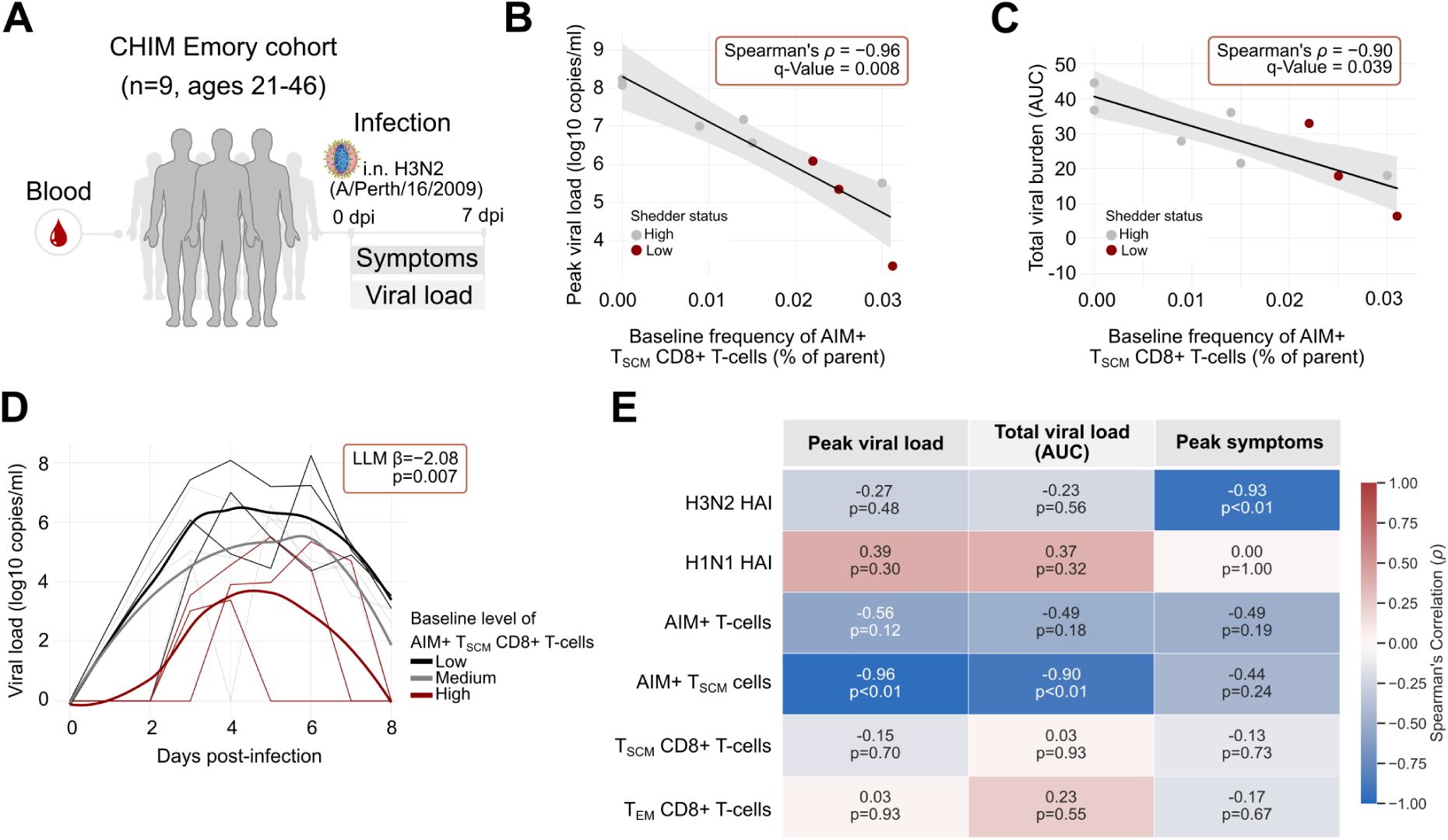
Association of baseline influenza-specific T_SCM_ cells with viral control in a human challenge model. **(A)** Schematic overview of the Controlled Human Infection Model (CHIM) Emory cohort study design. Nine healthy adults (ages 21-46) were challenged intranasally with influenza A/Perth/16/2009 (H3N2). Blood samples were collected at baseline (0 dpi). Symptoms and viral load was assessed for 7 days post-infection (dpi). **(B)** Spearman’s rank correlation (ρ=−0.96, q-Value = 0.008) between the peak viral load (log10 copies/ml) and the baseline frequency of AIM+ T_SCM_ CD8+ T cells (% of parent). **(C)** Spearman’s rank correlation (ρ=−0.90, q-Value = 0.039) between the total viral burden (Area Under the Curve [AUC]) and baseline AIM+ T_SCM_ frequency. **(D)** Viral load kinetics quantified from the viral RNA in nasopharyngeal swabs (log10 copies/ml) over 7 days post-infection. Individual trajectories are stratified by baseline AIM+ T_SCM_ CD8+ T cell levels: Low (black), Medium (grey), and High (red). Statistical significance of the longitudinal association was determined using a linear mixed model (LMM β=−2.08, p=0.007). **(E)** Correlation matrix heatmap displaying associations between baseline immune parameters (H3N2/H1N1 HAI titers, total AIM+ T cells, AIM+ T_SCM_ cells, T_SCM_ CD8+ T cells, T_EM_ CD8+ T cells) and clinical outcomes (Peak viral load, Total viral load AUC, Peak symptoms). Blue indicates a negative correlation and red indicates a positive correlation, with p-values listed within each cell.

We systematically tested a comprehensive matrix of baseline (pre-infection) immune features, including humoral immunity (H1N1 and H3N2 HAI) and a deep profile of T-cell subsets (including total and antigen-specific (AIM+) T_SCM_, T_EM_, and T_EMRA_ populations), against viral load and symptom severity following H3N2 infection (**Table S5, Fig. S5**). This analysis revealed a stark divergence in the mechanisms of protection.

The baseline frequency of influenza virus-reactive stem cell-like memory CD8+ T cells (AIM+ T_SCM_) emerged as the single strongest predictor of viral control. This specific population exhibited a robust and statistically significant negative correlation with peak viral load (Spearman’s ρ = -0.96, q = 0.008; **Fig. 6B, Fig. S5A,** *upper panel*) and total viral burden (AUC; Spearman’s ρ = -0.90, q = 0.039; **Fig. 6C, Fig. S5B,** *upper panel).* A longitudinal linear mixed-effects model (LMM) confirmed this finding, revealing that subjects with higher baseline AIM+ T_SCM_ levels experienced a more rapid viral clearance (**Fig. 6D**). The model identified a significant negative association between the baseline AIM+ T_SCM_ frequency and the viral load trajectory over time (β = -2.08, p = 0.007; **Fig. 6D**).

In striking contrast, AIM+ T-cell subsets, including AIM+ T_SCM_, showed no statistical association with symptom severity (ρ = -0.44, p = 0.24; **Fig. 6E, Fig. S5A, B,** *lower panels*). Instead, symptom mitigation was strongly and exclusively correlated with baseline humoral immunity. The H3N2 HAI titer was the most significant predictor of peak total symptoms (ρ=−0.93, q=0.065; **Fig. 6E**). This humoral correlate, however, had no significant association with viral load metrics (peak viral load ρ = -0.27, p = 0.48 and total viral load ρ = -0.23, p = 0.56; **Fig. 6E**).

Collectively, these findings from the human challenge study establish a functional bifurcation in protective immunity. Baseline humoral antibodies correlate exclusively with symptom mitigation, while the pre-existing frequency of influenza virus-reactive T_SCM_, the population mechanistically linked to our pediatric cohort, is the strongest correlate of viral control. The CHIM results serve as supportive, external validation consistent with the pediatric findings, highlighting the clinical importance of T_SCM_ for protective antiviral immunity.

## Discussion

The quest for a universal influenza vaccine is predicated on the ability to elicit broad, long-lasting immunity that transcends seasonal drift (*48*). While T stem cell-like memory (T_SCM_) cells have been identified as a theoretical reservoir for such durability (*6, 8–10, 12*), vaccine development has been hindered by an inability to expand this rare population in humans reliably. In this study, we identified the immunological setpoints that govern this process. We demonstrate that the expansion of influenza virus-reactive T_SCM_ cells is not a stochastic event but is strictly determined by a pre-existing state of innate antiviral readiness. Our systems immunology approach reveals that a baseline, pDC-associated type I IFN signature acts as the primary determinant of T-cell fate, directing responders toward a functionally poised, Th1-dominant phenotype while non-responders default to a dysfunctional, hyper-proliferative state. By validating these findings in a controlled human infection model, we further establish a clinical division of labor where T_SCM_ cells are responsible for rapid viral clearance, providing a mechanistic rationale for their prioritization in next-generation vaccine design.

Our work extends the paradigm of systems vaccinology beyond predicting peak immunogenicity to forecasting immunological durability. Seminal studies on the yellow fever (YF-17D) and seasonal influenza vaccines established that early post-vaccination signatures could predict peak antibody and effector T-cell magnitudes (*14–18, 49–52*). However, protection against recurrent threats requires not just an initial burst of effectors but the establishment of a self-renewing memory pool. By identifying a baseline signature that predicts the magnitude of the T_SCM_ compartment on day 28 after vaccination, we provide a proxy for long-term persistence. This is a critical conceptual advance, as the early establishment of T_SCM_ clones has been directly linked to the longevity of protection in chronic viral infections and mRNA vaccination (*53*). Furthermore, our validation in a human challenge cohort establishes a clear functional division of labor: pre-existing antibodies mitigate symptoms, while the baseline frequency of influenza virus-reactive T_SCM_ cells is the primary predictor of viral load control. This establishes the T_SCM_ subset as a distinct endpoint for next-generation vaccines aiming to limit viral transmission and disease severity in the absence of sterilizing humoral immunity.

A central and seemingly paradoxical finding of our study is that a baseline signature enriched for type I IFN predicts a positive vaccine outcome. This contrasts with the inflammaging literature where chronic baseline interferon signaling in the elderly is associated with immune exhaustion and refractoriness to vaccination (*54*). We propose that these observations represent two distinct biological states: tonic readiness and chronic exhaustion. The signature we identify in children, characterized by poised pDCs and elevated circulating IL-18 and CXCL10, likely reflects a low-and-ready state of innate training potentially maintained by recent subclinical exposures. Unlike the high-basal, refractory profile of inflammaging, which leads to receptor desensitization, this tonic state facilitates a rapid, coordinated response.

This beneficial state of innate readiness suggests a convergence with the concept of trained immunity, which describes the long-term functional reprogramming of innate immune cells through epigenetic modifications following a primary stimulus (*55–57*). This phenomenon mirrors the non-specific cross-protection observed with the Bacillus Calmette-Guérin vaccine, where BCG-induced remodeling of myeloid progenitors confers broad protection against heterologous respiratory viral infections (*58–60*). It is plausible that the tonic pDC signature identified in our responder cohort represents a form of naturally acquired trained immunity resulting from prior biological exposures. In this context, poised pDCs act as endogenous adjuvants that bridge the non-specific memory of the innate system with the specific memory of the adaptive system. This implies that strategies designed to induce trained immunity could be strategically leveraged to prime the host environment before influenza vaccination, thereby ensuring the optimal innate tone required for the optimal expansion of T_SCM_-mediated immunity (*59*).

Mechanistically, upon mucosal vaccination, we propose that these poised pDCs act as a trigger by releasing a decisive bolus of IFN-α. This provides the optimal third signal required to license APCs and guide T_SCM_-cell expansion away from terminal exhaustion and toward a more functionally poised fate. In responders, the antigen-specific T_SCM_ pool exhibited a functionally poised phenotype (Th1-high, Ki67-low), whereas non-responders generated cells characterized by proliferative stress and functional inertia. This suggests that the baseline innate environment does not merely gate the magnitude of the response but actively programs its quality. A robust and transient IFN signal appears to act as a Goldilocks stimulus, sufficient to drive productive expansion but regulated enough to allow cells to exit the cell cycle and arrest in a stem-like state. Conversely, the absence of this optimal priming in non-responders results in an abortive attempt at expansion, leading to a dysfunctional memory compartment. The validation of this pathway via structural equation modeling and the confirmation of the network via protein-protein interaction analysis centered on STAT1 and DDX58 move our findings from statistical correlation to a cohesive causal model.

The distinct clinical utility of the T_SCM_ compartment identified in our adult challenge model offers a compelling perspective on the long-term goal of pediatric vaccination. While the adult participants likely acquired their protective T_SCM_ repertoire through cumulative natural infections and various prior vaccinations rather than pediatric LAIV, the clear association between higher T_SCM_ levels and viral control in this group highlights the functional necessity of this compartment. In children, we observed that the magnitude of the T_SCM_ response was directly proportional to the number of prior LAIV doses. This suggests that repeated LAIV administration mimics natural exposure, progressively building a reservoir of durable cells similar to the protective profile observed in adults. By effectively seeding the long-lived T_SCM_ compartment during the plastic phase of the developing immune system, repeated live vaccination may therefore establish a foundational layer of cellular immunity, aiming to replicate the antiviral resilience typically acquired through natural infection, but without the risks of disease.

These findings have immediate implications for the design and evaluation of universal influenza vaccines. The historical reliance on hemagglutination inhibition titers has stalled the development of T-cell-based vaccines (*61*). Our identification of the T_SCM_ subset as a correlate of viral control supports the adoption of cellular endpoints in clinical trials. Moreover, our predictive signature offers a roadmap for precision vaccinology. The variability in baseline readiness suggests that a one-size-fits-all approach is inherently limited (*62*). We envision a predictive and corrective strategy in which a companion diagnostic based on the 32-gene ISG score could identify individuals with low innate readiness. These predicted non-responders could then be targeted with next-generation vaccines co-formulated with specific adjuvants, such as TLR7/9 or STING agonists encapsulated in lipid nanoparticles, designed to reconstitute the missing pDC-IFN axis artificially. The expansion of T_SCM_ against conserved epitopes in influenza viruses can also be combined with vaccination strategies aimed at inducing protective antibodies against subdominant conserved B-cell epitopes, as both strategies are not mutually exclusive.

Our study has limitations inherent to its design. The intense multi-omic profiling necessitated smaller cohort sizes. While the statistical robustness of our models and the biological consistency across cohorts provide confidence, larger studies will be required to refine the diagnostic thresholds for the IFN signature. Additionally, while our transcriptomic and proteomic data strongly implicate pDCs as the source of the innate signature, definitive confirmation would require depletion studies or single-cell analysis of tissue-resident populations, both of which are challenging in pediatric human cohorts. Finally, while we show that this innate state predicts T_SCM_ expansion, future interventional studies using specific adjuvants (e.g., TLR7/9 agonists) are needed to prove that inducing this state *de novo* can rescue non-responders.

In conclusion, our work provides a systems-level roadmap for the rational design of durable influenza vaccines. We move beyond the observation of heterogeneity to define the molecular mechanism, tonic innate readiness, that dictates the quality of T-cell memory. For clinical translation, this offers a two-fold application: first, as a predictive biomarker to stratify patients who may require boosted regimens or use of enhanced vaccines (i.e. high dose or adjuvant); and second, as a specific blueprint for next-generation adjuvants designed to mimic this pDC-associated IFN signature. By targeting the innate environment to expand the T_SCM_ reservoir deliberately, we can engineer the long-term viral control required for a truly universal influenza vaccine.

## Materials and Methods

### Study Design and Participants

#### Pediatric LAIV Cohort (FluPRINT)

We conducted a single-armed, phase 4 immunogenicity (*63, 64*)study (NCT04222595) in Oxford, UK, adhering to Good Clinical Practice standards with approval from the East Midlands, Nottingham 1 Research Ethics Committee and the UK Medical Research Council. Written informed consent was obtained from parents of all participants. The study enrolled healthy, immunocompetent children aged 4-6 years of European descent, excluding those with severe chronic conditions, autoimmune disorders, systemic immunosuppression, or recent live vaccine administration. This analysis focused on a sub-cohort of 19 participants stratified by prior LAIV history into naive (Group 1), one prior dose (Group 2), and two or more prior doses (Group 3). Whole blood samples were collected at baseline (day 0) and day 28 post-vaccination.

#### Controlled Human Influenza Infection Model (CHIM)

To validate clinical correlates of protection, we utilized samples from a single-center, open-label CHIM study conducted at Emory University (NCT05332899). The study enrolled healthy volunteers aged 18 to 49 years following rigorous screening to exclude chronic medical conditions. Under an Emory University IRB-approved protocol (STUDY00000083), participants were admitted to an inpatient quarantine unit and inoculated intranasally with influenza A/Perth/16/2009 (H3N2) (*63, 64*). Nasopharyngeal swabs were collected daily for 7 days post-inoculation to monitor viral shedding via qRT-PCR. Clinical outcomes were parameterized as Peak Viral Load (maximum viral copy number), Total Viral Burden (Area Under the Curve of the viral load trajectory), and a cumulative Symptom Score based on daily self-reporting using the influenza patient-reported outcome (FLU-PRO) diary (*65*). Peripheral blood was collected at baseline (pre-challenge) for immunological profiling.

### Sample Collection and Processing

For the FluPRINT cohort, whole blood collected at day 0 and day 28 was processed immediately. A 2 mL aliquot was stabilized in Paxgene tubes and stored at -80°C for transcriptomic analysis. The remaining volume was separated into plasma, which was stored at -80°C, and peripheral blood mononuclear cells (PBMCs), which were isolated via Lymphoprep density gradient centrifugation and cryopreserved in liquid nitrogen. For the CHIM cohort, PBMCs and plasma were isolated and cryopreserved using standard protocols compatible with the FluPRINT processing pipeline to ensure data comparability.

### Immunological Assays

#### Hemagglutination Inhibition (HAI) Assays

Serum HAI antibody titers were determined using standard protocols involving receptor-destroying enzyme treatment for 16-18 hours at 37°C followed by heat inactivation (*54, 55*). For the FluPRINT cohort, titers were assessed at day 0 and day 28 against A/Brisbane/02/2018 (H1N1), A/Kansas/14/2017 (H3N2), B/Colorado/06/2017 (Victoria), and B/Phuket/3073/2013. For the CHIM cohort, titers were determined at baseline against the specific challenge strain (A/Perth/16/2009 (H3N2)).

#### Mass Cytometry (CyTOF) and Antigen Stimulation

Thawed PBMCs were stimulated for 12-16 hours with a comprehensive peptide pool library spanning the entire proteome of influenza A/California/07/2009, consisting of 483 20-mers and 24 9-mers (*12*). Matched unstimulated controls were processed in parallel for every sample. Following stimulation, cells were stained with a 36-marker metal-labeled antibody panel designed to distinguish T-cell memory subsets and intracellular cytokine production (*12*). The workflow included surface labeling, fixation with Maxpar Fix I, permeabilization with Maxpar Perm Buffer, and intracellular staining. Data were acquired on a Fluidigm Helios mass cytometer and analyzed using Cytosplore software and FlowJo.

#### Targeted Plasma Proteomics

Plasma protein levels were quantified using the Olink Target 96 Inflammation panel (Olink Bioscience, Uppsala, Sweden), which measures 92 inflammation-related biomarkers via Proximity Extension Assay technology. Data were expressed as Normalized Protein Expression (NPX) values on a log2 scale. Quality control was performed using internal controls; samples were excluded if they deviated significantly from the control median. Missing values falling below the limit of detection were imputed using the median to preserve distribution centrality.

### Transcriptional Profiling and Bioinformatics

#### RNA Sequencing and Differential Expression

Total RNA was extracted from Paxgene-stabilized blood using the RNeasy Mini Kit, and libraries were prepared using the TruSeq Stranded Total RNA kit for sequencing on an Illumina HiSeq platform. Raw reads were aligned and quantified using Salmon. After filtering for low-abundance transcripts, differential expression analysis was performed using DESeq2 with a parametric fit. Differentially expressed genes were defined based on a log2 fold-change threshold of >0.3 and an adjusted p-value < 0.05.

#### Gene Set Variation Analysis (GSVA)

To infer biological process activity, we performed GSVA using the *GSVA* R package. We curated a library of gene sets from MSigDB, including Hallmark, C2 Curated, and C7 Immunologic collections. Additionally, we incorporated specific Viral Readiness signatures derived from published datasets (*29*) to quantify transcriptional responses to IAV, SARS-CoV-2, and RSV. Enrichment scores were calculated for each subject using log-transformed normalized counts.

#### *In Silico* Deconvolution

To determine the cellular composition of the bulk transcriptome, we employed *xCell* R package. Raw gene counts were converted to Transcripts Per Million and processed to generate enrichment scores for immune cell types. These scores were used to calculate specific composite cellular metrics, including the pDC Score (baseline pDC enrichment) and the APC-to-Lymphocyte Ratio.

### Computational Systems Immunology

#### Virtual Sorting of Antigen-Reactive T Cells

To rigorously identify rare influenza-specific CD8+ T cells, we developed a computational virtual sort pipeline. For each subject, the median expression intensity of markers in the matched unstimulated control was subtracted from the peptide-stimulated sample, with negative values clamped to zero. These background-subtracted data were normalized using robust scaling and subjected to dimensionality reduction via t-SNE and unsupervised clustering to define phenotypic metaclusters. We defined three functional metrics for AIM+ (CD69+CD137+) cells: Proliferative Stress (Ki-67 frequency), Th1 Score (median expression of IFNγ and TNFα), and a Th1/Proliferation Balance Score calculated as the difference between the Th1 and Proliferative Stress scores.

#### Responder Classification and Composite Scoring

We employed a data-driven Receiver Operating Characteristic (ROC) optimization to define T_SCM_ responders. We tested a range of post-vaccination T_SCM_ frequency thresholds and identified 1.4% as the optimal cutoff, maximizing the area under the curve for discriminating response magnitude. This binary classification was used for all group comparisons. To model the mechanistic cascade, we integrated transcriptomic and cellular data into standardized composite Z-scores representing the pDC Baseline, the ISG Score (mean of 32 canonical interferon-stimulated genes), the APC Priming Score (composite of antigen presentation genes and proteins), and the Th1 Response Score.

#### Multi-omic Data Integration

To model the determinants of the vaccine response at a systems level, we constructed a unified, multi-modal dataset integrating distinct biological layers for each subject (n=19). The final integrated matrix comprised 177 features spanning five categories: (1) Clinical Metadata (Age, Sex, Vaccination History); (2) Cellular Composition (Baseline frequencies of pDCs and macrophages derived from xCell deconvolution); (3) Transcriptional Signatures (Composite ISG Score); (4) Pathway Enrichment Scores (GSVA scores for key antiviral and metabolic pathways); and (5) Targeted Proteomics (Normalized Protein Expression values for inflammatory cytokines). Before integration, missing values were imputed using the median. To ensure comparability across modalities with differing dynamic ranges (e.g., gene counts vs. protein NPX), all continuous features were standardized (Z-score transformed) to a mean of 0 and standard deviation of 1. This unified matrix was utilized for all subsequent multivariate modeling (PLS-DA) and structural equation modeling (SEM).

#### Multivariate Modeling and Network Analysis

We assessed the variance explained by the global multi-omic signature using PERMANOVA with Euclidean distances. To test the hypothesized causal cascade, we constructed a Structural Equation Model (SEM) using the lavaan package with a Diagonally Weighted Least Squares estimator to handle mixed continuous and categorical data. Feature selection for biomarker discovery was performed using Partial Least Squares Discriminant Analysis (PLS-DA) to identify variables with high Variable Importance in Projection scores. Unsupervised consensus clustering was used to define data-driven patient immunotypes. Finally, to define the physical connectivity of the identified signatures, discriminative features were mapped to the STRING database (v12.0). Protein-protein interaction (PPI) networks were constructed using a high-confidence interaction threshold of 0.7, incorporating evidence from experiments, databases, text mining, and co-expression. Network topology was analyzed to identify central regulatory hubs based on node degree centrality.

### Statistical Analysis

All statistical analyses were performed in R and Python. Continuous variables were compared using the non-parametric Mann-Whitney U test for unpaired groups or the Wilcoxon signed-rank test for paired data. Multiple group comparisons used the Kruskal-Wallis test with Dunn’s post hoc correction. Correlations were assessed using Spearman’s rank coefficient. To identify the predictive plasma protein signature, we employed Lasso (Least Absolute Shrinkage and Selection Operator) regression with cross-validation to select the minimal robust feature set. For the CHIM cohort, associations between baseline immunity and viral kinetics were modeled using linear mixed-effects models with random subject intercepts. Predictive models were validated using Leave-One-Out Cross-Validation and permutation testing with 10,000 iterations. Statistical significance was defined as a two-tailed p-value < 0.05 or a False Discovery Rate (FDR) < 0.05.

## Data Availability

The RNA-sequencing data generated in this study have been deposited in the Gene Expression Omnibus (GEO) under accession code (GSE310641). Mass cytometry data are available via Zenodo (FluPRINT cohort: https://doi.org/10.5281/zenodo.17654309 and CHIM cohort: https://doi.org/10.5281/zenodo.17654451). The full integrative multi-omic dataset (19 subjects x 177 features) used for systems modeling is available via Zenodo (https://doi.org/10.5281/zenodo.17654510). The custom code and scripts used for the virtual sort, causal modeling, and systems analysis are available on GitHub at https://github.com/atomiclaboratory/fluprint_tscm. All other data supporting the findings of this study are available within the article and its Supplementary Information files or from the corresponding author upon reasonable request.

## Contributors

**A.T.** and **A.J.P.** conceived and designed the FluPRINT study. **A.T.** conceptualized the systems immunology approach, initiated the project, and wrote SOPs for sample acquisition for the FluPRINT cohort. **S.L.**, **N.R.**, and **M.D.P.** designed and conducted the Controlled Human Influenza Infection Model (CHIM) study. **E.C.**, **N.S.**, **P.A.**, **H.R.**, and **S.M.** managed clinical operations, laboratory work, and sample collection for the FluPRINT cohort. **D.A.** performed CyTOF and peptide stimulation experiments for both cohorts, under the supervision of **C.M. A.E.** and **T.A.** performed hemagglutination inhibition (HAI) assays for the FluPRINT cohort, under the supervision of **A.G.-S. I.T.** performed the primary data analysis, computational modeling, statistical testing, and generated all figures under the supervision of **A.T. J.T.** and **N.S.** conducted Gene Set Variation Analysis (GSVA). **S.H., M.F.,** and **H.N.** contributed to transcriptomic data processing. **A.T.** interpreted the data and wrote the original draft of the manuscript. All authors critically reviewed, edited, and approved the final version of the manuscript.

## Declaration of interests

AJP was previously the Chair of the UK Department of Health and Social Care’s Joint Committee on Vaccination and Immunisation and is a member of WHOs Product Development Advisory Committee. The A.G.-S. laboratory has received research support from Avimex, Dynavax, Pharmamar, 7Hills Pharma, ImmunityBio, and Accurius. A.G.-S. has consulting agreements for the following companies involving cash and/or stock: Castlevax, Amovir, Vivaldi Biosciences, Contrafect, 7Hills Pharma, Avimex, Pagoda, Accurius, Esperovax, Applied Biological Laboratories, Pharmamar, CureLab Oncology, CureLab Veterinary, Synairgen, Paratus, Pfizer, Virofend and Prosetta. A.G.-S. has been an invited speaker in meeting events organized by Seqirus, Janssen, Abbott, Astrazeneca and Novavax. A.G.-S. is inventor on patents and patent applications on the use of antivirals and vaccines for the treatment and prevention of virus infections and cancer, owned by the Icahn School of Medicine at Mount Sinai, New York, US. Other authors declare no competing interests.

## Acknowledgments

We express our deepest gratitude to the young study participants and their parents in the pediatric FluPRINT cohort, as well as the brave volunteers in the CHIM study who underwent controlled influenza infection. Without their participation and trust in our research endeavor, this work would not have been possible. We would also like to thank the dedicated team of nursing staff, the laboratory research support, and the clinical trials support offices at the Oxford Vaccine Group (OVG) at the University of Oxford. We extend our appreciation to the research assistants at OVG for collecting samples at odd hours. Special thanks to Amy Beveridge for her invaluable assistance with laboratory logistics and general support. We appreciate helpful discussions and support from all members of the OVG. We thank the Oxford Genomics Centre at the Wellcome Centre for Human Genetics (funded by Wellcome Trust grant reference 203141/Z/16/Z) for generating and initial processing the sequencing data, with special thanks to Angela Lee. This work was partly supported by CRIPT (Center for Research on Influenza Pathogenesis and Transmission), a NIAID-funded Center of Excellence for Influenza Research and Response (CEIRR, contract # 75N93021C00014) to AGS and TA. This study was funded by the EC Marie Curie Fellowship award (granted to AT; FluPRINT, Project 796636), the MRC Hic-VAC award to AT, and the NIHR BRC to AJP. The CHIM study at Emory University has been supported, in whole or in part, by Flu Lab.

## Members of the CHIM Study Group

Anice Lowen, Department of Microbiology and Immunology, Emory University; Jessica Traenkner, Hope Clinic, Emory University; Cecilia Zhang, Hope Clinic, Emory University; Meredith J. Shephard, Department of Microbiology and Immunology, Emory University; Michelle N. Vu, Department of Microbiology and Immunology, Emory University; A.J. Campbell, Department of Microbiology and Immunology, Emory University; Shamika Danzy, Department of Microbiology and Immunology, Emory University; Nahara Vargas-Maldonado, Department of Microbiology and Immunology, Emory University.

## Supplementary figures

**Figure S1.**
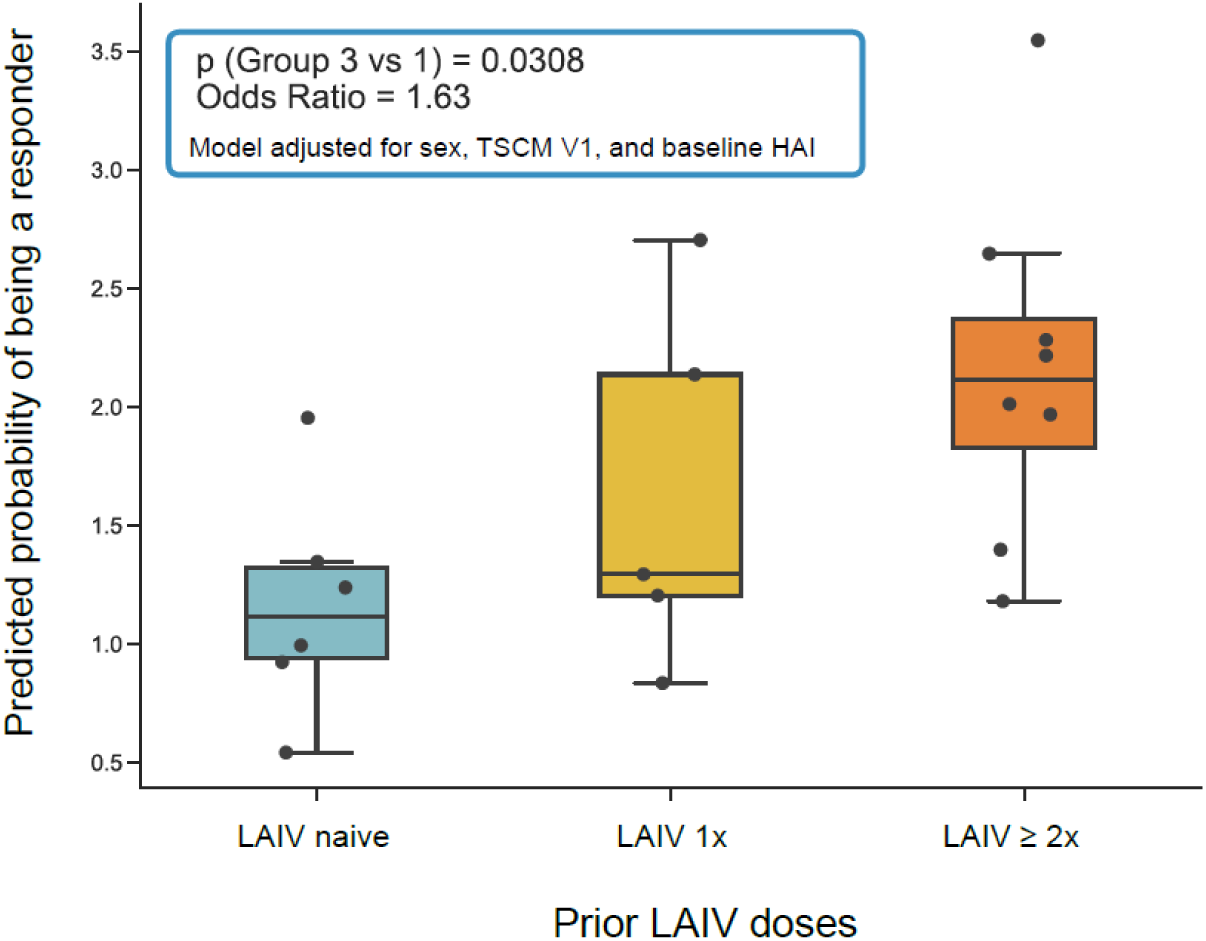
Predicted probability of being a responder based on prior LAIV exposure. Box plots display the predicted probability of classification as a responder (Y-axis) stratified by the number of prior Live Attenuated Influenza Vaccine (LAIV) doses received: Naive (0 doses), 1 dose, and ≥2 doses. The logistic regression model was adjusted for sex, baseline antigen-specific stem cell-like memory T cell (TSCM) levels (Visit 1), and baseline Hemagglutination Inhibition (HAI) titers. The comparison between the high-exposure group (≥2x) and the naive group was statistically significant (p=0.0308), with an odds ratio of 1.63.

**Figure S2.**
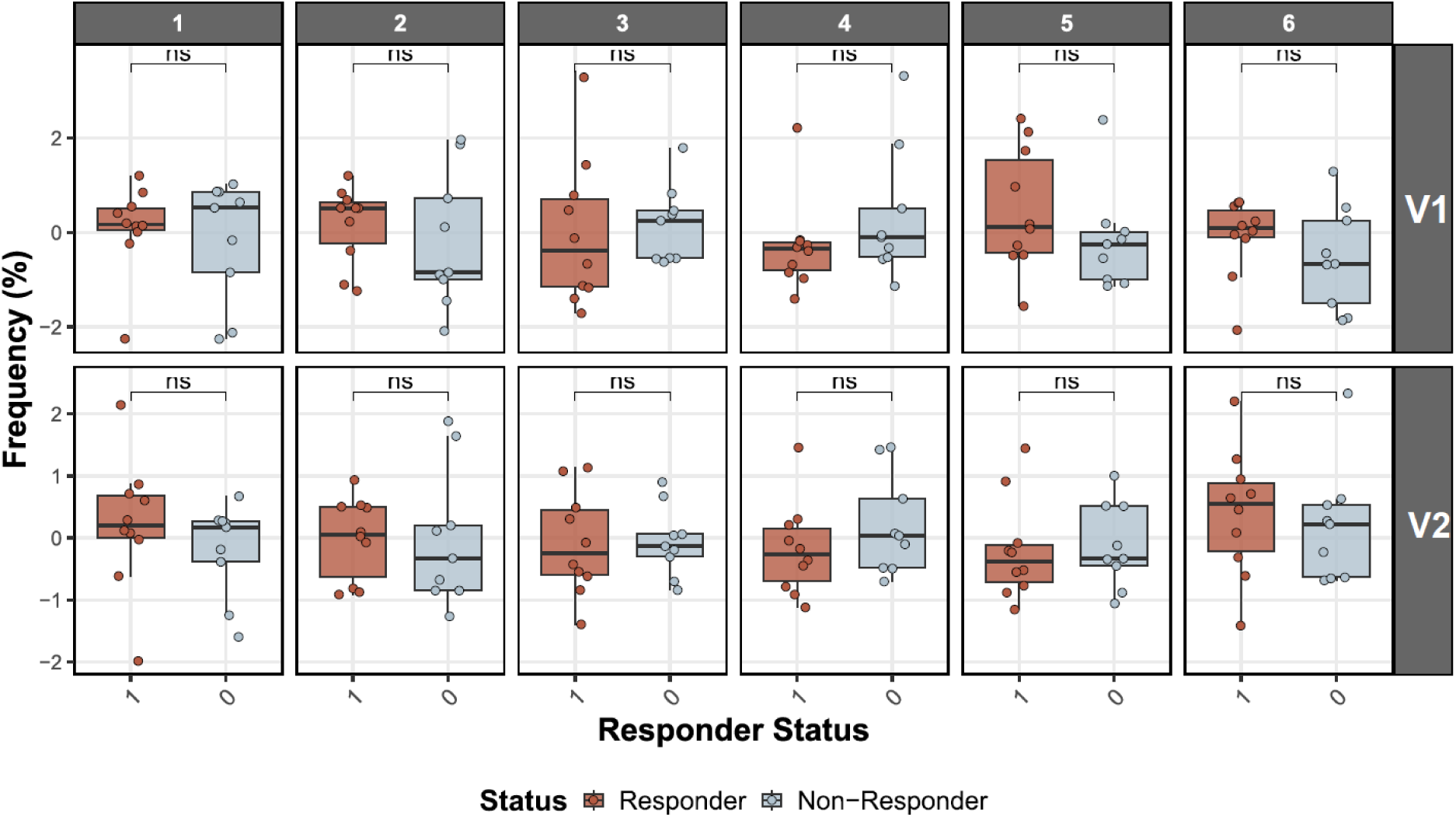
Frequency of immune cell clusters by responder status and time point. Box plots showing the frequency (%) of six distinct immune cell clusters (Columns 1-6) stratified by responder status. (*Top row*) Cluster frequencies at baseline (Visit 1; V1). (*Bottom row*) Cluster frequencies post-vaccination (Visit 2; V2). Responders are shown in red, and non-responders are shown in blue. Significance testing indicated no statistically significant differences (“ns”) between groups for these clusters at the measured time points.

**Figure S3.**
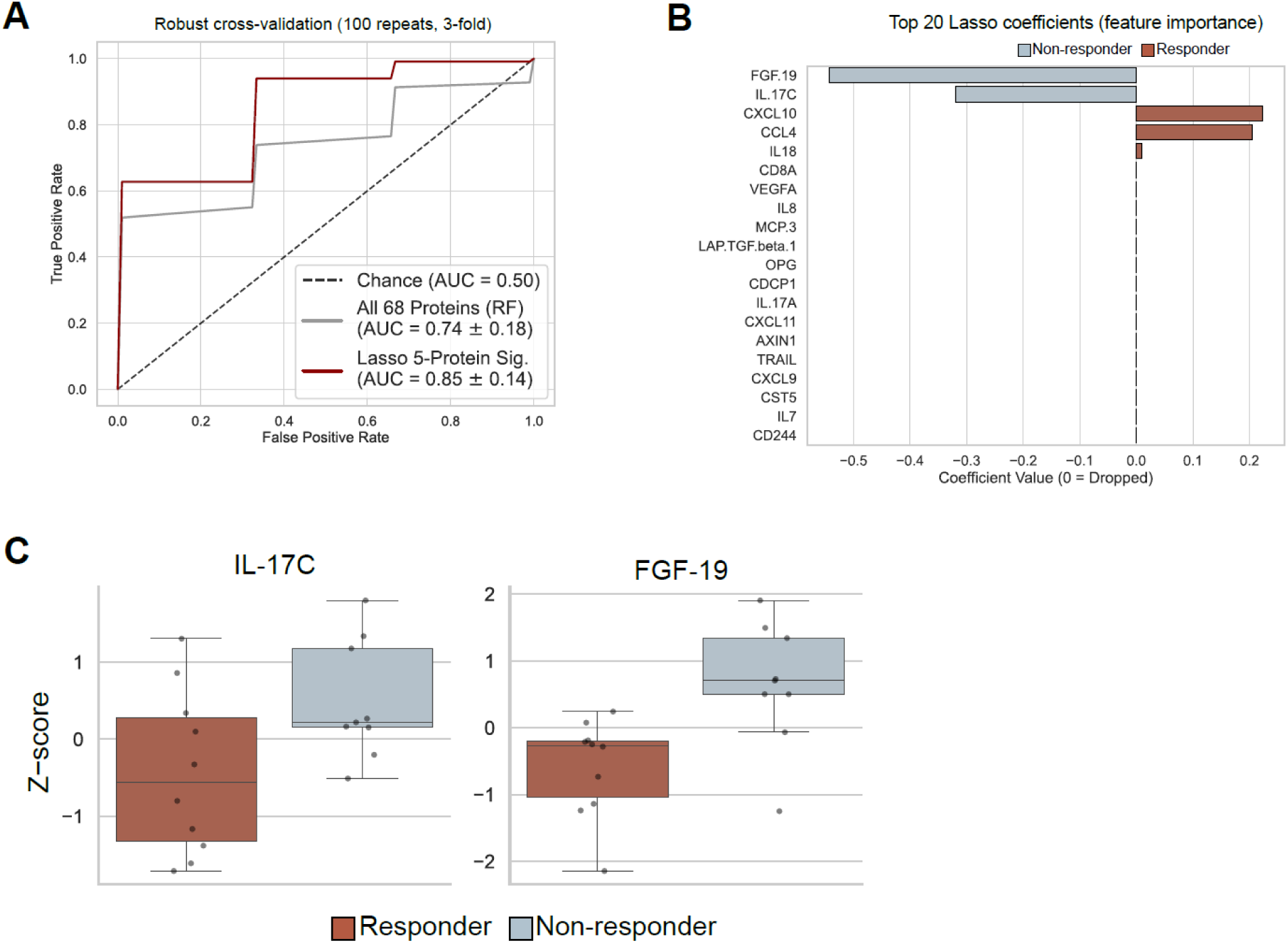
Predictive modeling of responder status using plasma protein signatures. **(A)** Receiver Operating Characteristic (ROC) curves comparing the performance of classifiers trained to predict responder status. The grey line represents a Random Forest model trained on all 68 measured proteins (AUC = 0.74), while the red line represents a Lasso regression model trained on a refined 5-protein signature (AUC = 0.85). Performance was evaluated using robust cross-validation (100 repeats, 3-fold). **(B)** Bar plot displaying the top 20 Lasso coefficients (feature importance). Bars extending to the right (red) represent features positively associated with responders (CXCL10, CCL4, IL18), while bars extending to the left (blue) represent features associated with non-responders (FGF-19 and IL-17C). **(C)** Box plots comparing the normalized Z-scores of the top negative predictors, IL-17C and FGF-19, between responders (red) and non-responders (blue).

**Figure S4.**
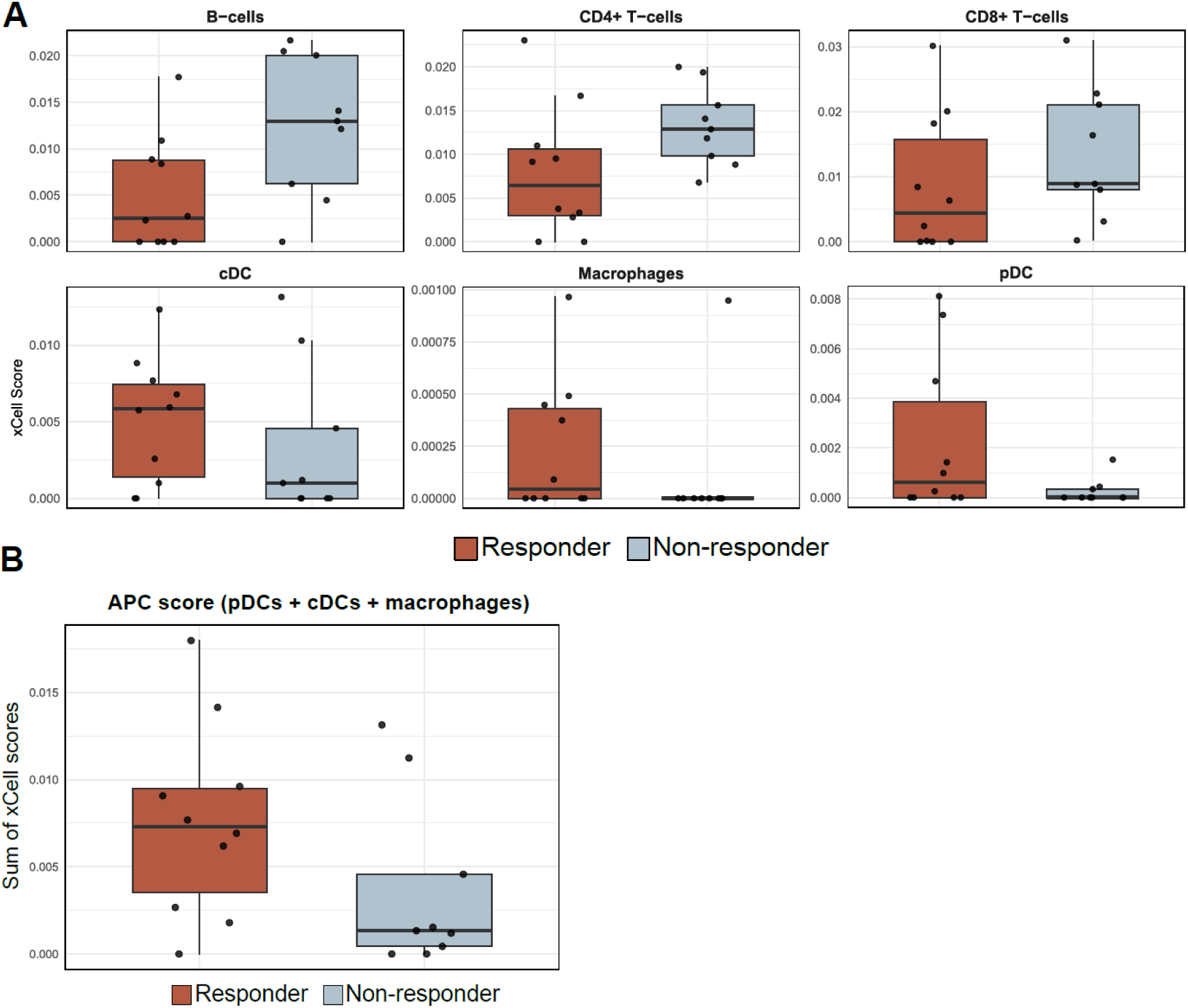
Comparison of predicted cell type abundance between responders and non-responders. **(A)** Box plots displaying the cell enrichment scores (xCell Score) for specific immune subsets, B-cells, CD4+ T-cells, CD8+ T-cells, cDCs, Macrophages, and pDCs, stratified by responder status (Responder: red; Non-responder: blue). **(B)** Composite Antigen Presenting Cell (APC) score, calculated as the sum of xCell scores for pDCs, cDCs, and macrophages, comparing responders vs. non-responders.

**Figure S5.**
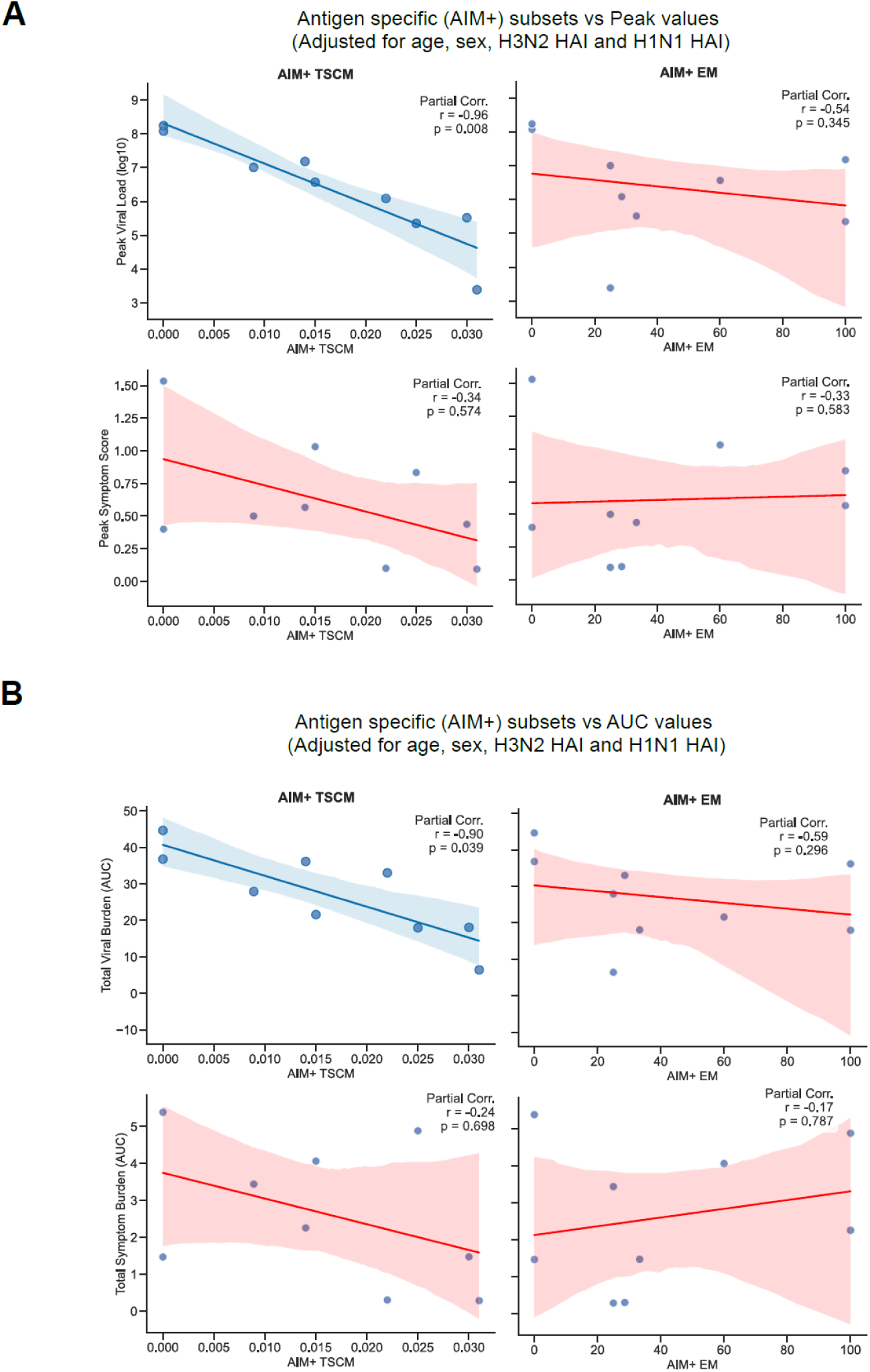
Correlations between antigen-specific T-cell subsets and clinical outcomes. Scatter plots with regression lines showing the partial correlations between baseline Antigen-Specific (AIM+) TSCM subsets and clinical outcomes. All analyses were adjusted for age, sex, and baseline H3N2 and H1N1 HAI titers. **(A)** Correlation of AIM+ TSCM levels with peak viral load (top) and peak symptom score (*bottom*). **(B)** Correlation of AIM+ TSCM levels with total viral burden (Area Under the Curve, AUC; *top*) and total symptom burden (AUC; *bottom*). Partial correlation coefficients (r) and p-values are displayed within each panel. Shaded areas represent 95% confidence intervals.

## Supplementary tables

**Table S1. Causal mediation analysis results.** This analysis estimates the Average Causal Mediation Effects (ACME) and Average Direct Effects (ADE) across three different model specifications (n=19). The results determine the proportion of the total effect mediated through the hypothesized causal pathways.

**Table S2. Path analysis results connecting pDC baseline to responder status.** This analysis evaluates the sequential relationships between pDC baseline levels, ISG scores, APC priming scores, and Th1 response scores in predicting responder status (n=19).

**Table S3. Path analysis results connecting macrophage baseline to responder status.** This analysis evaluates the sequential relationships between macrophage baseline levels, ISG scores, APC priming scores, and Th1 response scores in predicting responder status (n=19).

**Table S4. Feature loadings for Component 1.** This table presents the loading values for the top features driving Component 1. Features include specific genes, pathway database signa-tures, and derived clusters. Negative and positive loadings indicate the direction of association with the component.

**Table S5. Correlation analysis between immune features and clinical outcomes**. This table summarizes the Spearman rank correlations (ρ) between specific immune features (T-cell sub-sets and HAI titers) and clinical outcomes (Viral Load and Symptom scores). N = 9 observations per comparison.

## Notes

### Competing Interest Statement

A.J.P. was previously the Chair of the UK Department of Health and Social Care's Joint Committee on Vaccination and Immunisation and is a member of WHOs Product Development Advisory Committee. The A.G.-S. laboratory has received research support from Avimex, Dynavax, Pharmamar, 7Hills Pharma, ImmunityBio, and Accurius. A.G.-S. has consulting agreements for the following companies involving cash and/or stock: Castlevax, Amovir, Vivaldi Biosciences, Contrafect, 7Hills Pharma, Avimex, Pagoda, Accurius, Esperovax, Applied Biological Laboratories, Pharmamar, CureLab Oncology, CureLab Veterinary, Synairgen, Paratus, Pfizer, Virofend and Prosetta. A.G.-S. has been an invited speaker in meeting events organized by Seqirus, Janssen, Abbott, Astrazeneca and Novavax. A.G.-S. is inventor on patents and patent applications on the use of antivirals and vaccines for the treatment and prevention of virus infections and cancer, owned by the Icahn School of Medicine at Mount Sinai, New York, US. Other authors declare no competing interests.

